# Cophylogeny simulators are not interchangeable: similarities, differences and structural biases in synthetic host-symbiont data

**DOI:** 10.64898/2026.09.13.751162

**Authors:** Gabriele Di Palma, Catherine Matias, Blerina Sinaimeri

## Abstract

Synthetic data are becoming increasingly important for computational studies of cophylogeny, including machine learning inference, benchmarking, and method testing. Several generators have been proposed to produce such data, but each relies on different assumptions about host-symbiont coevolution. These assumptions are often implicit and rarely examined, even though results can depend strongly on the synthetic model being used. In this article, we present a systematic structural analysis of representative cophylogeny generators under controlled scenarios. The goal is to make their assumptions explicit and to understand how these choices shape the synthetic data they produce as well as the conclusions that may be drawn from them.

## 1. Introduction

Reconstructing the shared evolutionary history of symbionts and their hosts is central in several application domains, including the identification and tracking of emerging infectious diseases (Etherington et al., 2006; Lei and Olival, 2014; Pennington et al., 2015). The growing availability of public sequence data has made these analyses increasingly feasible at scale. A standard way to formalize host-symbiont coevolution is through *cophylogeny* models, which aim to explain the histories of hosts and symbionts using their evolutionary trees (typically inferred from DNA sequences). In this framework, coevolution is often expressed as the problem of mapping the symbiont phylogeny onto the host phylogeny (see, e.g., Charleston, 2003; Donati et al., 2015; Menet et al., 2022; Page, 1994). This mapping, called a *reconciliation*, describes the relationship between the two trees via biologically meaningful events such as cospeciation, host-switching, duplication, and loss. Because real host-symbiont datasets are limited in number, size, and diversity, synthetic data are an important resource for developing and validating computational methods in cophylogeny. Several synthetic generators for host-symbiont systems have been proposed, based on different modeling assumptions and simulation strategies. These generators are used to evaluate reconciliation methods and statistical tests of congruence, and they are expected to become even more important for data-driven pipelines, especially machine learning, where large, controlled datasets are needed for training and benchmarking. Despite this growing role, synthetic generators are often treated as interchangeable, and their outputs are rarely analyzed in detail beyond the specific task they are used for. In practice, different generators may encode different assumptions about evolutionary processes and may constrain the space of host-symbiont structures they can produce. As a result, datasets generated under comparable evolutionary scenarios may still differ substantially in their trees structures, detectable coevolutionary signal, and association network topology. Such differences can directly affect the conclusions drawn from synthetic-based computational studies.

In this article, we study the structural properties of synthetic host-symbiont datasets produced by commonly used cophylogeny generators. Thus, we adopt a regime-based evaluation framework, considering cospeciation-dominated, host-switch-dominated, and mixed coevolutionary regimes, and analyze each simulated dataset at three complementary levels. **First**, we examine coevolution-based summaries of the generated data, focusing on the relative frequencies of key events (in particular cospeciation and host-switching) to verify how closely each generator follows the intended regime. **Second**, we compare the generators outputs relying on two types of characteristics. On one hand, tree-based measures characterize the host and symbiont phylogenies through size and shape descriptors, including the number of leaves and standard tree-balance indices such as Cherry, Sackin, and Colless. On the other hand, association-based measures characterize the bipartite network defined by the host-symbiont nowadays associations. We measure classical network parameters such as the connectivity and heterogeneity (e.g., density, assortativity, and hosts hotspot concentration). **Third**, we study the effects of a generator choice on a downstream analysis such as parsimonious reconciliation. Finally, we compare these structural characteristics of the synthetic data to those observed in real host-symbiont datasets, which we use more as a reference set rather than as ground truth. This study is guided by three questions:

> *Q*_1_ When we fix the same high-level coevolutionary regime, do different generators produce datasets with comparable structure, or do they systematically induce different tree shapes, congruence signals, and association-network patterns?
>
> *Q*_2_ Do differences among synthetic generators propagate to the output of a standard downstream cophylogenetic analysis, namely parsimonious reconciliation?
>
> *Q*_3_ How much do the structural patterns produced by each generator overlap with those observed in real host-symbiont datasets, and do the generators fully cover this structural reference space or leave significant regions occupied by real data unexplored?

Our goal is not to identify a “best” generator but to explore how different modeling choices determine the structure of synthetic cophylogenetic data. By making these differences clear, we aim to help researchers choose synthetic data in a careful way for cophylogeny studies, and to point out the biases that can come from using one generator instead of another.

## 2. Methods

### 2.1. Description of the evaluated generators

We focus on the host-symbiont setting, where the generators simulate a host phylogeny, a symbiont phylogeny, and the associations between them. Existing generators in the literature follow different modeling choices (for example in the set of macro-evolutionary events they include, or whether they use time and branch lengths). We therefore compare the following host-symbiont generators, namely `Coala` (Baudet et al., 2014), `treeducken` (Dismukes and Heath, 2021) and the model described in Alcala et al. (2017) that has no name and that we call `cophylo` (from the name of the main function provided by the authors). We also include `AsymmeTree` (Schaller et al., 2022), originally developed in the gene-species setting. Although its gene-family simulation builds on ideas from tools such as `SaGePhy` (Kundu and Bansal, 2019), `AsymmeTree` provides a simpler “tree evolving on a tree” framework that produces reconciliation-style histories without introducing additional layers such as domain evolution or population processes. This makes it structurally closer to host-symbiont generators and allows a more direct comparison in our setting. We do not consider gene-species simulators more broadly, since many of them include processes specific to gene evolution, such as horizontal gene transfer with replacement or incomplete lineage sorting, which do not have clear counterparts in host-symbiont macro-evolutionary models. Conversely, host-symbiont systems may involve *multiple associations*, where a symbiont is linked to more than one host at a time, a situation that does not arise in the gene-species setting.

We now describe the four generators considered in this study. Notice that most of these generators are by-products of more global coevolution or cophylogenetics methods that we do not describe here. We rather only focus on the parts of these tools that enable generating a pair of host and symbiont trees together with coevolution history and extant leaves associations. Some of these tools only generate a symbiont tree *conditional* on a given input host tree (and additional parameters), while other *jointly* generate the host and symbiont tree (in general, from a birth and death process). In order to get a full generating process of host and symbiont pair, we combine the former tools with a simple birth-death process with parameters (*λ*_*H*_, *µ*_*H*_) (speciation and extinction, respectively) on the host tree when it should be given as input.

#### 2.1.1. Coala

The `Java` software `Coala` (Baudet et al., 2014; Sinaimeri et al., 2023) contains a generator (`TGLGenerator.jar`) for simulating symbiont coevolution along a given host tree and allowing four different types of events: *cospeciation, duplication, host-switch* and *loss*. The input host tree does not need to be dated (if branch lengths exist, they will be discarded) so we rely on the function `TreeSim` from the `DendroPy python` library (Moreno et al., 2024a,b), using parameters (*λ*_*H*_, *µ*_*H*_) and conditioned on the number of extant leaves to produce the input host tree. The output symbiont tree will similarly be undated. The generator uses three parameters resulting in a vector ⟨*p*_*c*_, *p*_*d*_, *p*_*s*_, *p*_*l*_⟩ of four probabilities constrained to sum to 1. The model is based on the Duplication-Transfer-Loss (DTL) model (Bansal et al., 2012; Tofigh et al., 2011). Conditional on a given host tree *H*, the generation of the symbiont tree proceeds by recursively considering its unmapped nodes that are (temporarily) “positioned” on branches and for which one of the four different events occurs. The process starts by positioning the symbiont root on the branch before the root of *H* (by default, although the user may choose any host branch), where this initial branch is fictitious and serves only to initialize the simulation. For every unmapped symbiont node that is temporarily positioned on a host branch, the three events cospeciation, duplication and switch (so except in case of a loss) will induce its mapping on the host node below the branch it was positioned on, as well as its speciation. Now, in case of a cospeciation (occurring with probability *p*_*c*_), each descendant symbiont lineage is positioned on one of the two host branches below the associated host. Conversely, when duplication occurs (with probability *p*_*d*_), each descendant symbiont lineage stays positioned on the same host branch its parent was on. A host-switch (occurring with probability *p*_*s*_) induces one descendant lineage to be positioned on the same host branch its parent was on, while the other descendant is positioned on a host branch chosen randomly, under the constraint that this does not violate the time feasibility of the reconstruction so far (Stolzer et al., 2012). If such a branch does not exist, the switch does not happen and a new event *i* is drawn among the three others (namely *i* ∈ *{c, d, l}*) with rescaled probability *p*_*i*_ */*(*p*_*c*_ + *p*_*d*_ + *p*_*l*_). Finally when a loss occurs (with probability *p*_*l*_ = 1 − *p*_*c*_ − *p*_*d*_ − *p*_*s*_), the symbiont node is not mapped and rather re-positioned on one of the 2 host branches (chosen randomly) below the branch it was. Note that a loss event can represent three distinct yet indistinguishable scenarios: (a) speciation of the host species independently of the symbiont, which then remains associated with only one of the new host species; (b) cospeciation of the host and symbiont, “immediately” followed by the extinction of one of the newly formed symbiont species; or (c) the same as (b), but with a failure to detect the symbiont in one of the two new host species. The process continues until no unmapped symbiont nodes remain; the evolution of a lineage stops once it is mapped to a host leaf. For a fixed host tree *H* and a given set of event probabilities, the simulator generates multiple symbiont trees (100 by default). A representative tree is then selected as the median tree, *i.e*., the tree that minimises the distance to the others under a chosen tree-distance measure. Note that the set of events included in `Coala` result in host-symbiont coevolved trees with no multiple associations between their leaves (so that a symbiont has at most one host). The method `AmoCoala` (Sinaimeri et al., 2023) later developed to overcome this limitation relies on an initial pair of host-symbiont trees and their leaves associations to set additional local probabilities of new events (called *spreads*) that produce these multiple associations and are defined at every internal node of the host tree. This very-fine tuning of the model (allowing heterogeneous probabilities of spreads along the tree) becomes a drawback in our study, as the sheer number of additional choices would complicate our analysis. For this reason, we choose not to include this generator in our comparison.

#### 2.1.2. treeducken

The `sim_cophyBD` function from the `R` package `treeducken` (Dismukes and Heath, 2021) implements a cophylogenetic birth-death process that simulates, in forward time, a pair of phylogenies (host tree *H* and symbiont tree *S*) together with their extant ecological interactions (Dismukes and Heath, 2021). Note that it is the only tool out of the four tested here that simulates the two trees jointly. The model allows hosts and symbionts to undergo speciation and extinction independently, and it also includes coupled events such as *cospeciation* (with explicit parameter) and *coextinction* (implicit).

The generator is based on a continuous-time Markov process and is conditioned on a simulation time horizon. Both trees will start at the same date and evolve during that time horizon, creating trees with the same stem age. The generator contains a total of six competing events, parameterized by their rates (*λ*_*H*_, *µ*_*H*_, *λ*_*S*_, *µ*_*S*_, *χ, λ*_*c*_). Symbiont evolution is governed by the speciation rate *λ*_*S*_ and the extinction rate *µ*_*S*_ . A symbiont speciation event is similar to a duplication-like event from `Coala`, in which both descendant lineages inherit the ancestral host associations, while a symbiont extinction corresponds to a (particular case of a) loss-like event in that setting. The model includes a *host-expansion* event that takes two different forms (*switch* and *spread*), both being (additional) symbiont speciation events occurring with rate *χ*. The switch event creates one symbiont speciation with one descendant that has a new host association (among hosts not associated to its parent) while the other descendant retains the ancestral host repertoire. In spread event, one randomly chosen descendant gains a novel host association in addition to the ancestral host repertoire (thus creating a multiple association). The tool has three options (accessible through `hs_mode` ∈ {`“switch”,”spread”,”both”}`), where only switches, spreads or both events (with half probability for each) may occur, respectively. The user may also enforce a *host limit, i.e*., a maximum number of host associations per symbiont lineage at any given time. Cospeciation occurs at rate *λ*_*c*_ when a host lineage bifurcates and induces a simultaneous symbiont speciation event, triggered on one randomly selected symbiont lineage currently associated with that host. After cospeciation, each descendant host lineage is associated with one of the two new symbiont descendants, and remaining ancestral associations are redistributed at random among descendants. Hosts can also speciate independently of symbionts at rate *λ*_*H*_ and any associated symbiont will then remain associated to each descendant (thus creating a multiple association). This is analogous to a failure-to-diverge event as described in Charleston (2009) and implemented in Jane (Conow et al., 2010). Finally, host extinction occurs at rate *µ*_*H*_ and may implicitly induce coextinction of symbiont lineages that are left without any associated hosts.

#### 2.1.3. cophylo

The article Alcala et al. (2017) comes with a code as Supporting Information (Alcala et al., 2017). It includes a `C` program (`main_cophylo.c`) generator for simulating a symbiont tree *S*, conditional on the input host *H*, using both a continuous-time Markov process with three competing events: *speciation, extinction* and *host shift*; as well as an additional process of *cospeciation* that may occur at each host speciation. The input host should be a dated tree and we simulate it using the auxiliary function `species_tree_n_age` from the tool `AsymmeTree` (described below), that generates a birth-death process, conditional on the number of extant leaves but also on a time horizon *T* . Then the conditional simulation of the symbiont tree is obtained by successively running the former Markov process on each time interval defined by two successive host speciations. The generator has five evolutionary parameters: *λ*_*S*_, *µ*_*S*_ and *s* are the symbiont speciation, extinction and switch rates, respectively. Additionally, *c* is a cospeciation probability and *T*_*MRCA*_ a time to the most recent common ancestor (TMRCA). It enables to start simulating the symbiont tree at a different age than the host tree.

The first symbiont is linked to a random number *d* (drawn from parameter distribution *f*_*d*_ on the number of hosts per symbiont at any time) of contemporary hosts, or all available hosts if *d* exceeds their number. From the initial time to the next host speciation event, or between any two successive host speciation events, a continuous-time Markov process with the three above mentioned competing events is run. With rate *λ* (resp. *µ*) times the number of existing symbionts, a speciation (resp. extinction) event occurs to one symbiont randomly chosen. In case of a speciation, a first descendant inherits all associations of its parent while the other inherits a random number *d* (drawn with *f*_*d*_) among a subsample of these. In case of an extinction, that symbiont is simply removed. With rate *s* times the total number of current associations, a switch event occurs, randomly chosen among symbionts weighted by their number of current hosts. This symbiont gains a new host among those not yet associated to it. Note that in that switch event, contrarily to what happens in `Coala` or `treeducken`, the symbiont does not undergo a speciation. So that this event creates a multiple association. It rather corresponds to the spread event from `treeducken`. Additionally, at each host speciation time, their associated symbionts may cospeciate with probability *c*, in which case two symbiont descendants are created and each one follows one of the two descendant host. If there is no cospeciation, then the symbiont may either be a *specialist* (with probability *f*_*d*_ (1) which is user-chosen or ≈ 0.46 by default) or a *generalist*. A specialist will follow only one of the two descendant hosts, while a generalist will follow both and thus create a new multiple association. Note that in the latter case, the symbiont did not speciate and remains the same in the two descendant hosts, differentiating this case from the cospeciation one. The tool also admits a parameter for the minimum number of symbiont leaves, redoing the simulation whenever this number is not attained.

#### 2.1.4. AsymmeTree

The article Schaller et al. (2022) contains a general model for gene family history simulation along a species tree with *duplication, loss, horizontal transfer and conversion*. Putting aside the conversion event that is specific to the gene-species context, the model is similar to DTL mentioned above and may be used in the host-symbiont context. In the corresponding `python` package (Schaller et al., 2022), we focus here on two functions, namely `species_tree_n_age` that uses a birth-death model conditioned on time and number of extant leaves for simulating a species tree (host tree in our context), and `dated_gene_tree` that simulates a gene tree (symbiont tree in our context), conditional on a dated species tree. Note that in the former, the branches leading to extinct species are pruned from the output. We use the latter with only three events: duplication with rate *λ*_*D*_, loss with rate *λ*_*L*_ and horizontal (gene) transfer with rate *λ*_*HGT*_ . The root of the gene (symbiont) tree is placed at the root of the species (host) tree. The host-speciation events are ordered and processed sequentially. The simulation alternates between two cases: a host-speciation event and random events occurring in the symbiont tree during two successive host speciation times. When host-speciation occurs, the gene necessarily “cospeciates” at the same time; in our host-symbiont case, that means the symbiont cospeciates with its host with probability 1. Then, and similarly to `cophylo`, between two successive host-speciations, a continuous-time Markov process is run with the three competing events: duplication, loss and horizontal transfer. Duplication is the same event as in `Coala`, corresponding to symbiont speciation in `treeducken` or `cophylo`. A loss event, is the same as a loss in `Coala` or as a symbiont extinction in `treeducken` or `cophylo`. Finally, horizontal transfer corresponds to a switch in `Coala` or `treeducken` (and does not have an equivalent event in `cophylo`, since it implies a symbiont speciation). Note that this generator never produces multiple associations (that do not exist in the gene-species context).

#### 2.1.5. Summary of the generators characteristics

Table 1 summarizes the characteristics of the tools with a focus on the different coevolutionary events, while Table 2 contrasts these characteristics by emphasizing joint or conditional simulation of the trees as well as the role of time. Note that in the former, the event “Induced symbiont extinction” only exists for `treeducken` because the host and symbiont trees are jointly simulated, while for the other tools, the host tree is generated and pruned for extinct branches before being input in the conditional generator for the symbiont tree.

**Table 1.**
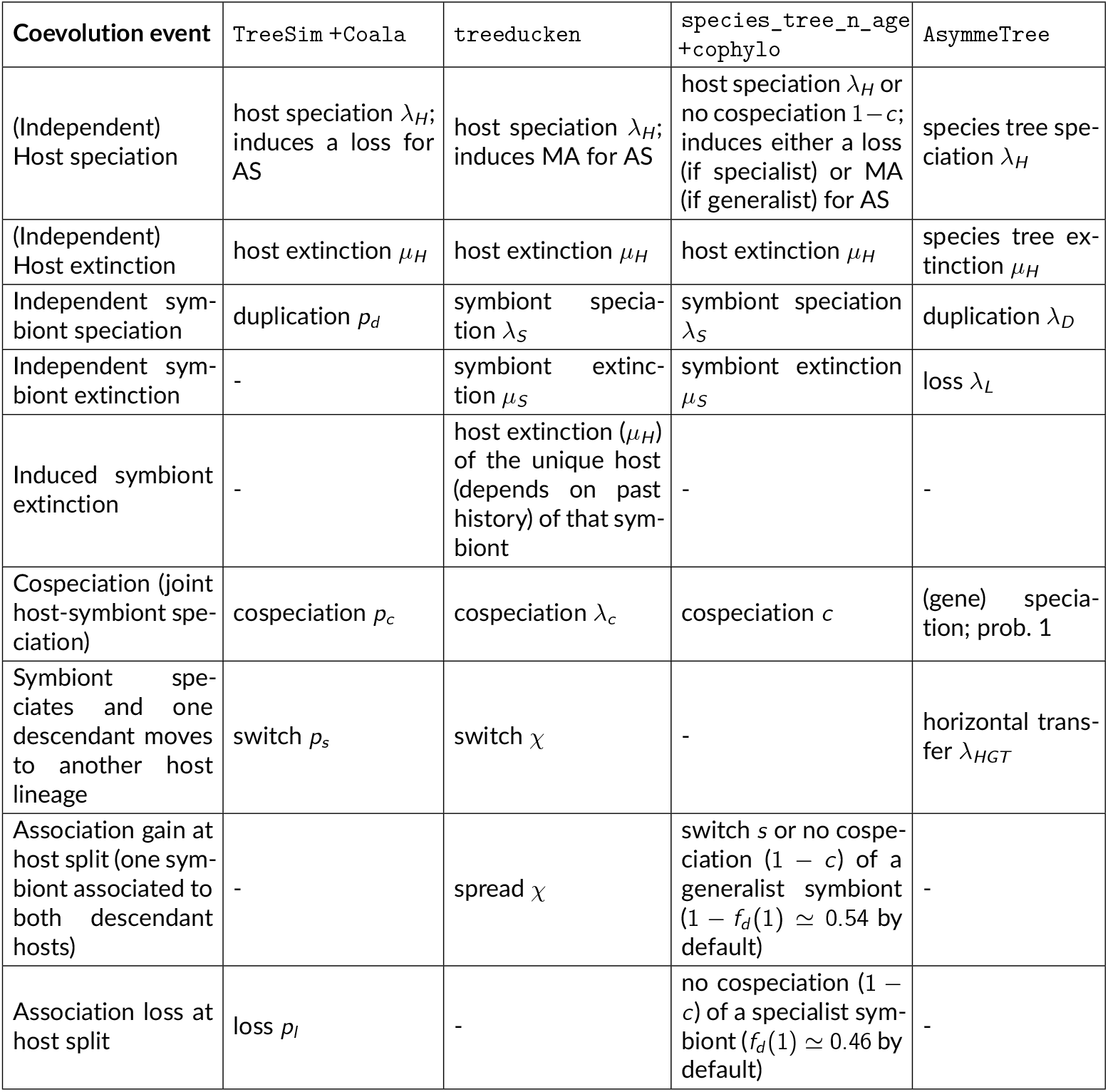
Correspondence between biological processes, event names and parameters across models. AS stands for associated symbionts; MA for multiple associations.

**Table 2.** Comparison of simulation design across cophylogeny generators.

| Model | Host phylogeny | Symbiont phylogeny | Associations | Time | Symbiont root |
| --- | --- | --- | --- | --- | --- |
| Coala | fixed input (simulated independently) | conditionally simulated | conditionally simulated | no | mapped to a host node |
| treeducken | jointly simulated | jointly simulated | jointly simulated | yes | same origin as host |
| cophylo | fixed input (simulated independently) | conditionally simulated | conditionally simulated with constraint on nb of hosts per symbiont at any time (input possible) | yes | independent origin time |
| AsymmeTree | birth-death process | conditionally simulated | conditionally simulated | yes | same origin as host |

To make the simulation settings as comparable as possible, we align host-generation parameters whenever they have a direct counterpart across methods, including the host speciation rate and, for `Coala`, `cophylo` and `AsymmeTree`, the host extinction rate within each regime (see section 2.2 for details). However, the methods still rely on different host-generation procedures. Since the topology of the host tree influences the resulting symbiont phylogeny, the effects of host and symbiont generation cannot be fully disentangled. We therefore interpret our results as comparisons of the complete simulation pipelines under the considered configurations. This reflects the comparison that a user can realistically make when applying these tools in practice, since their different host-generation mechanisms cannot be made fully identical without altering the tools themselves.

### 2.2. Coevolutionary regimes calibration and simulation design

We considered three main coevolutionary scenarios: a cospeciation-dominated regime *R*_1_, a host-switch-dominated regime *R*_2_, and a mixed regime *R*_3_. To make the intended scenarios explicit, we formalize them as regions in the plane of observed event frequencies. For each simulated dataset *i*, let *c*_*i*_ and *s*_*i*_ denote its observed cospeciation and host-switch frequencies, respectively. We focus on these two events because they are present, with comparable interpretation, across all generators. Notice that since *c* and *s* are frequencies computed with respect to the same total number of events, it should hold *c* + *s* ≤ 1. We define

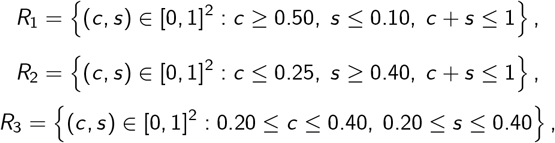

as illustrated in Figure 1.

**Figure 1.**
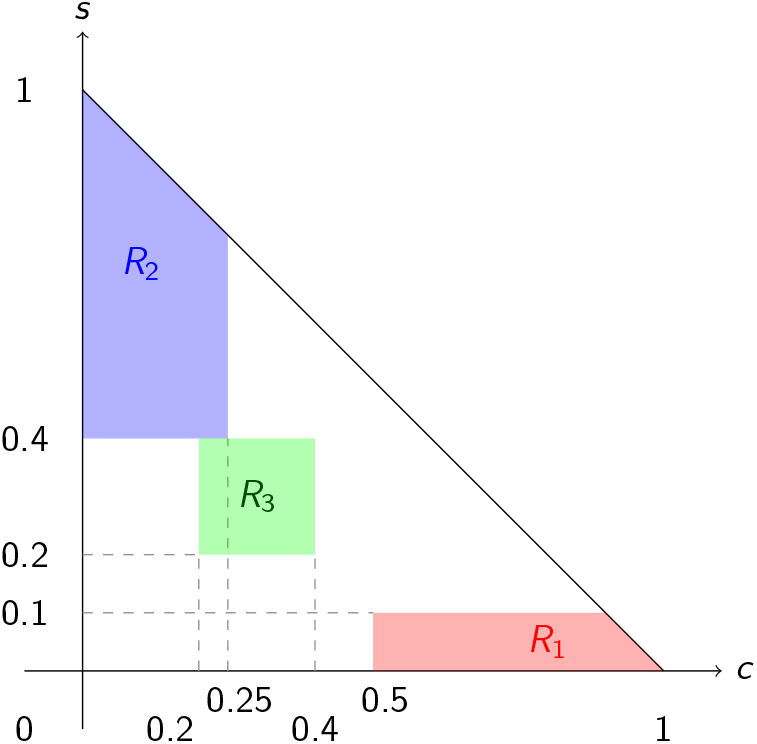
The 3 coevolution regimes. Cospeciation (*c* on *x* -axis) and host-switch (*s* on *y* - axis) frequencies (among all events) satisfy *c* +*s* ≤ 1. *R*_1_ region is cospeciation dominated, *R*_2_ is host-switch dominated, and *R*_3_ is a mixed regime.

For each generator-regime combination, we simulated 1000 synthetic datasets, denoted *D*_*G,q*_ with *G* ∈ {`Coala`, `treeducken`, `cophylo`, `AsymmeTree}` and *q* ∈ {1, 2, 3}. For each intended regime, generator-specific parameters were used to best approximate the scenarios, as reported in Table 3. More precisely, after defining the regions above in terms of the frequencies that we wanted to observe in the final datasets, we selected the input parameters of each generator so as to favour those outcomes. The numerical values of the input parameters do not coincide with the boundaries of the regions because only part of the simulated events are cospeciations or host switches. For example, to obtain a final history in which cospeciation represents at least half of all events, the cospeciation parameter may need to be set above 0.50, since duplications, losses, and host switches also occur and reduce its observed proportion. Moreover, the four generators control these events in different ways: `Coala` and `cophylo` use explicit parameters, whereas `treeducken` and `AsymmeTree` use rates that interact with the rates of the other events.

**Table 3.**
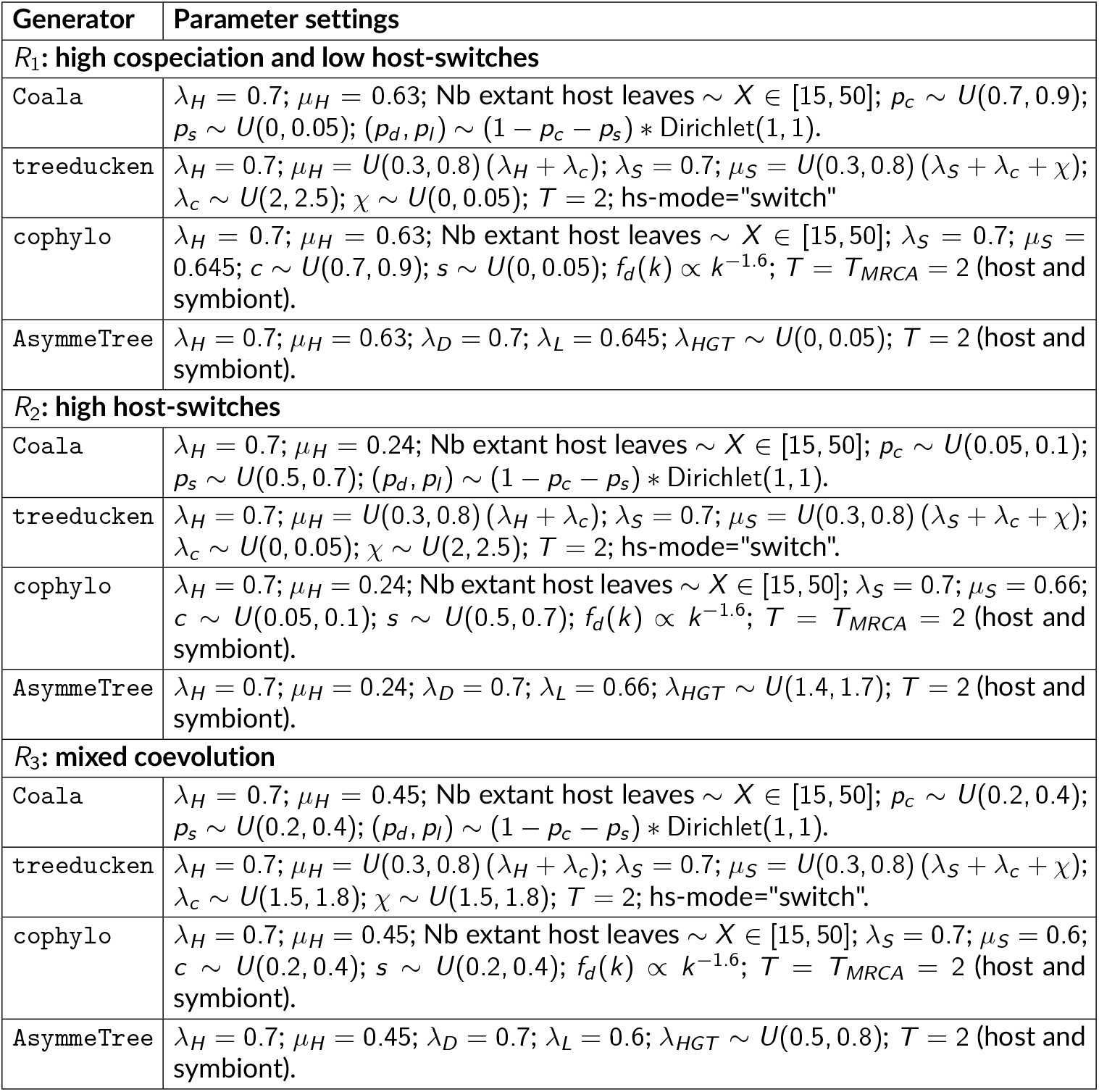
Parameter settings for regimes *R*_1_-*R*_3_ across generators.

The parameters were therefore chosen separately for each generator to produce datasets as close as possible to the same output regions.

We constrained the number of host leaves to range between 15 and 50 in all methods but `treeducken`, where host-tree size is an outcome of the joint simulation process. In order to reduce the number of parameters in `treeducken`, we used a parametrization proposed in Zeng and Román-Palacios (2025) where extinction rate *µ*_*H*_ (resp. *µ*_*S*_) is proportional to a total growth rate *λ*_*H*_ + *λ*_*C*_ (resp. *λ*_*S*_ + *λ*_*c*_ + *χ*). To make the tools as comparable as possible, we choose to set up the time *T*_*MRCA*_ in `cophylo` equal to the horizon time used to generate the symbiont tree. As a consequence, the host and the symbiont trees have the same age (as in the two other tools where time is considered). We also need to set up the distribution *f*_*d*_ on the number of hosts per symbiont at any time for that tool. Following the suggestion of the authors on their own dataset example, we rely on a power law with exponent parameter *α* = 1.6 and support in {1, …, |*L*(*H*)|} where *L*(*H*) is the number of extant hosts (*i.e* the probability to have *k* hosts is proportional to *k*^−*α*^). The minimum number of symbiont leaves in `cophylo` was set up to 5.

### 2.3. Evaluation of regimes reachability

For each generator *G* and intended regime *q* with *q* ∈ {1, 2, 3}, we quantify how often *R*_*q*_ is reached by the generator by looking at the following proportion, called *attainment proportion*,

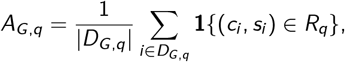

where |*D*_*G,q*_| is the cardinality of the corresponding set of simulated datasets (so 1000 in our study, see subsection 2.2), and **1**{(*c*_*i*_, *s*_*i*_) ∈ *R*_*q*_} is an indicator function equal to 1 if dataset *i* falls within region *R*_*q*_, and 0 otherwise. Moreover, to quantify how far the generated datasets are from the intended regime, we calculate the Euclidean distance of each observed dataset from the region corresponding to the regime:

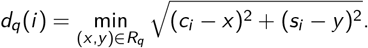

Thus, *d*_*q*_(*i*) = 0 when dataset *i* falls inside the intended region *R*_*q*_, whereas positive values indicate how far it lies outside the region.

The two measures provide different information. For example, two generators may have the same attainment proportion, while the datasets falling outside *R*_*q*_ are much closer to the region for one generator than for the other. Conversely, two generators may have similar distance summaries but different attainment proportions, meaning that the target region is reached more often by one of them. Considering both measures therefore gives a more complete picture of how well a generator reproduces the intended regime.

For each generator and regime, we report the attainment proportion *A*_*G,q*_, the median and interquartile range (IQR) of the values *d*_*q*_(*i*), and the observed medians of cospeciation and host-switch frequencies. Together, these quantities measure how closely the simulated datasets reproduce the intended coevolutionary scenario.

### 2.4. Measures used to compare the generators

To address *Q*_1_, we describe two different type of characteristics that we have measured on each pair of host and symbiont trees, across the different generators.

#### 2.4.1. Tree-based measures

For each simulated pair of host and symbiont trees, we first record basic size and scale characteristics. All distances and heights reported in those measures are defined in the topological sense (number of edges), rather than using branch lengths. This is because not all generators produce comparable temporal information, and some models enforce a fixed time horizon while others allow simulated times to extend beyond it. Thus, using topological distances ensures that the measures are comparable across generators.

For each tree (host or symbiont), we measure its number of leaves and its height (*i.e*. the distance from the root to its deepest leaf). For each simulated pair, we also compute the difference |*L*(*H*)| − |*L*(*S*)| between the number of leaves in the host and symbiont tree. We then compute standard shape and balance indices, through normalized versions of the *Cherry, Colless* (Colless, 1982) and *Sackin* (Sackin, 1972) indices, as follows.

For any rooted binary tree *T*, let *L*(*T*) and *V* (*T*) denote its sets of leaves and vertices, respectively.

A *cherry* is a pair of leaves that are adjacent to a common ancestor node. The *normalized Cherry index Ch*(*T*) of a tree *T* is defined as its number of cherries divided by the maximal possible value ⌊|*L*(*T*)|*/*2⌋ . The *normalized Colless index* (Colless, 1982) is defined as

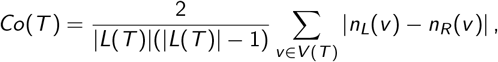

where *n*_*L*_(*v*) and *n*_*R*_ (*v*) denote the number of descendant leaves in the left and right subtrees of node *v* . The *normalized Sackin index* (Sackin, 1972) is defined as

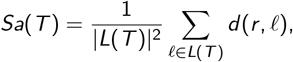

where *d* (*r, ℓ*) is the distance from the root *r* to leaf *ℓ*.

#### 2.4.2. Association-based measures

For each simulated dataset, we consider the bipartite host-symbiont interaction network defined on the leaves of the two trees. On this interaction network, we measure the *density*, the *degree-assortativity* and the frequency of *host hotspots*, as follows.

For a pair of host-symbiont trees (*H, S*), let *A* ⊆ *L*(*H*) × *L*(*S*) denote the set of associations, where (*h, s*) ∈ *A* if symbiont *s* is associated with host *h*. We write *G* = (*L*(*H*) ∪ *L*(*S*), *A*) for the resulting bipartite graph, which corresponds to the ecological interaction network of interest. The *density* of *G* is defined as

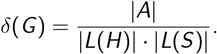

It is the fraction of possible connections that are present in the ecological network between hosts and symbionts.

Let deg(*v*) denote the degree of a node *v* in *G* . The *degree assortativity* (see Eq. (21) in Newman, 2003) measures the tendency of nodes to connect to other nodes with similar degree. It is defined as

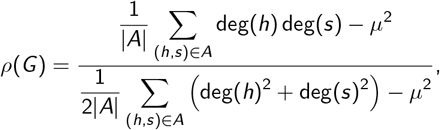

where

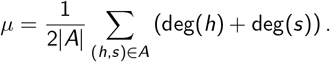

It measures the tendency of generalist species to be associated with other generalists, and specialists with other specialists. Positive values indicate assortative mixing, whereas negative values indicate that highly connected species tend to interact with less connected species.

A *host hotspot* is defined as a host leaf *h* ∈ *L*(*H*) whose degree exceeds the average host degree plus one standard deviation, a standard threshold used to identify highly connected nodes in ecological and interaction networks. Formally, letting

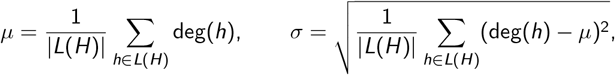

we define the set of host hotspots as

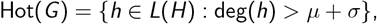

and record the frequency |Hot(*G*)|*/*|*L*(*H*)|. The frequency of hosts hotspots captures the presence of hosts that are highly connected (Newman, 2018; Poulin, 2010).

### 2.5. Downstream analysis: parsimonious reconciliation

To study the downstream effect of the choice of a generator, we used the classical event-based parsimony framework for cophylogenetic reconciliation. Given a host tree, a symbiont tree, and the associations between their extant leaves, a reconciliation explains the symbiont history in terms of four events: cospeciation (*C*), duplication (*D*), host-switch (*S*), and loss (*L*). A cost is pre-assigned to each event, and the cost of a reconciliation is the sum of the costs of all its events. For a given cost vector ⟨*w*_*C*_, *w*_*D*_, *w*_*S*_, *w*_*L*_⟩, a parsimonious reconciliation is therefore a reconciliation of minimum total cost (Charleston, 2003, 1998).

#### 2.5.1. Data filtering

The classical four-event model assumes that each symbiont leaf is associated with a single host leaf. It therefore does not directly represent multiple associations, where the same symbiont may be associated with several host species. Some reconciliation methods, such as `Jane` (Conow et al., 2010), allow particular forms of multiple association, but only through restricted mechanisms, such as failure of a symbiont to diverge. More general situations in which a symbiont spreads to an additional host require additional evolutionary events and are not represented by the classical four-event model (Sinaimeri et al., 2023). We therefore restricted the downstream analysis to datasets that could be converted into a *valid* input for the classical single-host-association reconciliation model, namely two binary phylogenetic trees together with leaf associations in which each symbiont is associated exactly with one host. Importantly, we did not simply discard every dataset containing multiple associations. Instead, for each synthetic dataset when a symbiont leaf was associated with several host leaves, we randomly kept one host association and removed the others. Host leaves unused by the retained associations (if any) were then pruned, since these are not normally part of the input considered by classical parsimonious reconciliation methods. Finally, we required both resulting trees to contain at least two leaves. Datasets for which this procedure produced fewer than two leaves in either tree, or an invalid tree structure were filtered and were recorded as failures.

Moreover, for computational feasibility, we imposed a maximum running time on each reconciliation analysis. Runs that did not terminate within this time limit were excluded from the subsequent profile and ambiguity analyses. Such cases typically corresponded to datasets with a very large number of optimal event vectors and optimal reconciliations. Since computational complexity depends on the reconciliation cost vector, the number of successfully analysed datasets may differ between the two cost settings (see section 3.3.3 below).

#### 2.5.2. Choice of event costs

The set of optimal reconciliations depends on the costs assigned to the four events. We considered the two classical cost vectors ⟨*w*_*C*_, *w*_*D*_, *w*_*S*_, *w*_*L*_⟩ = ⟨0, 1, 2, 1⟩ and ⟨0, 1, 3, 1⟩. The same two cost vectors were used for all generators and all regimes. We did not adapt the costs to the event frequencies of each generator. Our goal is not to recover the generating history, but to compare the outputs obtained when datasets from different generators are analysed with the same parsimonious reconciliation procedure.

#### 2.5.3. Optimal reconciliations and event vectors

An important point is that a minimum parsimony cost does not in general identify a unique reconciliation. Several different reconciliations may have exactly the same minimum cost, and these reconciliations may also differ in their numbers of coevolutionary events. Selecting arbitrarily one optimal reconciliation would therefore retain only one of the possible optimal event compositions and make the subsequent comparison across generators depend on an arbitrary choice.

To avoid making such an arbitrary choice, we used the tool `Capybara` (Wang et al., 2020). `Capybara` allows us to consider the different event compositions attained by optimal reconciliations. It associates to each optimal reconciliation its *event vector* (*N*_*C*_, *N*_*D*_, *N*_*S*_, *N*_*L*_), where the four entries are the numbers of cospeciation, duplication, host-switch, and loss events, respectively.

For each dataset and cost vector, we considered all distinct event vectors attaining the minimum parsimony cost. Furthermore, to compare datasets of different sizes, for each optimal event vector we computed the observed cospeciation and host-switch frequencies, namely

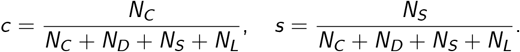

Thus, each optimal event vector is represented by a point (*c, s*) that we call *optimal event profile*.

#### 2.5.4. Comparison across generators

To address *Q*_2_, we compared generators separately for each regime and each cost vector. For each valid dataset *i*, we considered all *m*_*i*_ distinct optimal event vectors and mapped each of them to its corresponding optimal event profile (*c, s*).

To obtain one summary point for each dataset and cost vector, we computed the mean of the profiles associated with its distinct optimal event vectors:

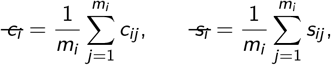

where *m*_*i*_ is the number of distinct optimal event vectors for dataset *i*.

#### 2.5.5. Number of optimal event profiles

Different optimal event vectors may lead to the same optimal event profile (*c, s*). Therefore, in addition to the dataset-level mean profile, we measured for each dataset and cost vector the number of distinct optimal event profiles,

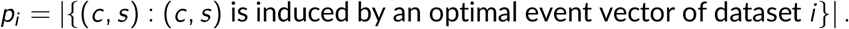

This quantity measures the ambiguity of the parsimonious analysis with respect to the relative frequencies of cospeciation and host-switching. We compared the distribution of *p*_*i*_ across generators separately for each regime and cost vector.

### 2.6. Comparison with real datasets

In order to address *Q*_3_, we considered an ensemble ℛ of 30 real host-symbiont datasets from the literature. These datasets span a broad range of biological systems, including plant-fungus interactions, insect-parasite and vertebrate-parasite associations, as well as endosymbiotic relationships (e.g., *Wolbachia*), and display substantial variation in tree sizes and association patterns (see Table 4).

**Table 4.**
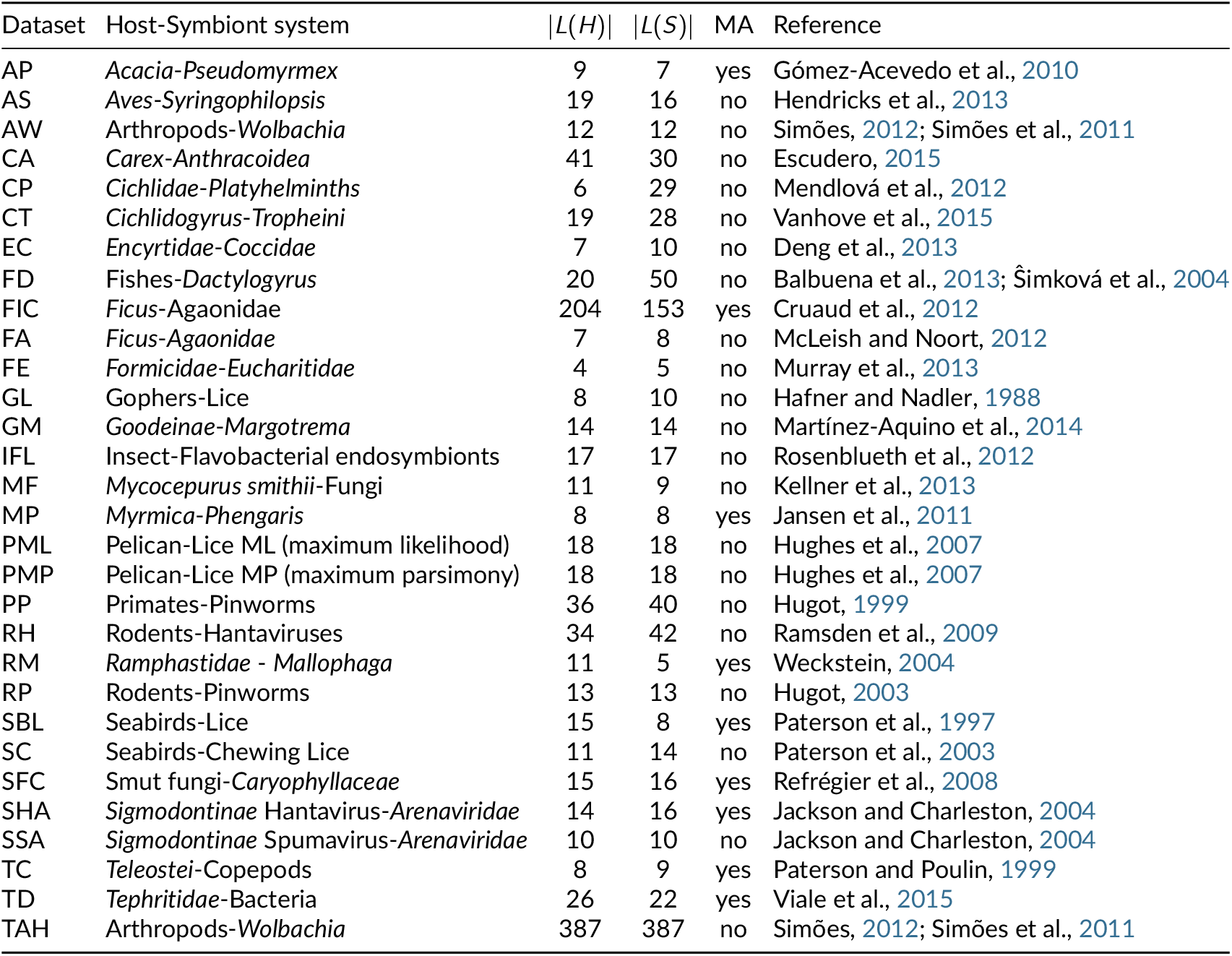
Host-symbiont datasets forming the reference set ℛ. For each dataset, we report the host-symbiont system, the number of host leaves |*L*(*H*)|, the number of symbiont leaves |*L*(*S*)|, and whether multiple associations (MA) are present.

It is important to note that the real datasets do not come with regime labels (e.g., “cospeciation-dominated” vs “switch-dominated”), so we cannot assign each real dataset to a specific simulated setting. Instead, we compared the real datasets separately with the synthetic datasets generated by each generator under each regime setting.

#### 2.6.1. Features choice

We relied on features that can be computed for both real and synthetic datasets. Each dataset *i* with (*H*_*i*_, *S*_*i*_) as host-symbiont pair of trees was represented by

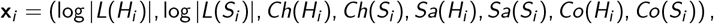

where |*L*(*H*_*i*_)| and |*L*(*S*_*i*_)| are the numbers of host and symbiont leaves, respectively, and *Ch, Sa*, and *Co* are the normalized Cherry, Sackin, and Colless indices (see section 2.4.1). We used the logarithm of the numbers of leaves to reduce the effect of very large trees.

Each dataset was represented by the eight features above. Before the comparison, each feature was standardized using its mean and standard deviation across the 30 real datasets, giving rise to standardized vector **z**_*i*_ . Thus, the standardized values describe each dataset relative to the empirical mean and variability of that feature.

#### 2.6.2. PCA visualization

We fitted PCA on the standardized features of the real datasets and projected the synthetic datasets onto the same first two principal components. PCA was used only to visualize the position of real and synthetic datasets in the same feature space. All quantitative comparisons described below were computed using the full set of eight standardized features.

#### 2.6.3. Proximity to real datasets

For the quantitative comparison, we used the full set of eight standardized features, rather than the PCA projection. For each synthetic dataset *s* ∈ *D*_*G,q*_, we computed its Euclidean distance to each real dataset and retained the smallest value

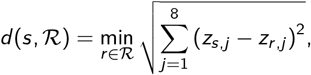

where we recall that ℛ is the set of real datasets and *z*_*i,j*_ is the standardized value of feature *j* for dataset *i* . A small value means that the synthetic dataset is close to at least one real dataset. For each generator and regime (*G, R*_*q*_), we report the median and IQR of these distances (among the 1000 synthetic datasets from *D*_*G,q*_).

#### 2.6.4. Coverage of the real data

The previous measure asks whether synthetic datasets from *D*_*G,q*_ are close to some real dataset in ℛ, but it does not show whether the different real datasets are represented by the synthetic data. We therefore also considered the comparison in the opposite direction. So for each real dataset *r* ∈ ℛ, we computed

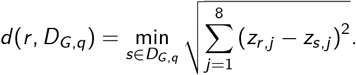

Small values reflect that the real dataset has a similar synthetic dataset. For each generator and regime, we report the median and maximum of these distances (among the 30 real datasets). The maximum identifies the real dataset that is least well represented by synthetic data.

#### 2.6.5. Proximity and coverage rates

The distance summaries describe how far synthetic and real datasets are from each other, but they do not directly show how frequently a generator produces datasets close to the reference data or how much of the reference dataset collection it represents. We therefore complemented the distances with two percentages. The first gives the proportion of synthetic datasets that are close to at least one real dataset, while the second gives the proportion of real datasets that have at least one close synthetic counterpart. To compute these percentages, we needed a reference value for what should be considered a small distance. We defined this reference from the distances observed among the real datasets themselves.

For each real dataset, we computed its distance to the closest other real dataset. We then set *τ* to the 95th percentile of these distances. For the 30 real datasets considered here, this gave *τ* = 1.945.

For each combination of a generator *G* and a regime *R*_*q*_, we compute the *proximity rate* Prox_*G,q*_ as the percentage of synthetic datasets *s* ∈ *D*_*G,q*_ for which *d* (*s*, ℛ) ≤ *τ* . It therefore gives the percentage of synthetic datasets that are within the empirical reference distance of at least one real dataset.

Similarly, we define the *coverage rate* Cover_*G,q*_ as the percentage of real datasets *r* ∈ ℛ for which *d* (*r, D*_*G,q*_) ≤ *τ* . It corresponds to the percentage of real datasets that have at least one synthetic dataset within the same reference distance.

#### 2.6.6. Feature-level comparison

Finally, we examined which individual features differ between real and synthetic datasets. For each generator, regime, and feature, we computed the median standardized value across the corresponding synthetic datasets and displayed these values in a heatmap. Since the features were standardized using the real datasets, a value of zero corresponds to the empirical mean. Positive values indicate values above the empirical mean, while negative values indicate values below it.

## 3. Results and discussions

### 3.1. Generators ability to reach coevolutionary regimes

Figure 2 shows the observed cospeciation and host-switch frequencies for the four generators under each intended regime. The shaded areas represent the three target regions. As many points may overlap in that representation, Figure 3 displays the boxplots for these same values. Table 5 reports the proportion of datasets inside each region and corresponding summary statistics. Finally, Figure 4 shows boxplots of the distance of each dataset from its target region.

**Table 5.** Agreement between the simulated datasets and the intended coevolutionary regimes. Attainment proportion *A*_*G,q*_ is the percentage of datasets from generator *G* whose observed cospeciation and host-switch frequencies fall within the corresponding region *R*_*q*_. The last column reports the median distance from the region, with the interquartile range in parentheses.

| Regime | Generator | $A_{G,q}$ (%) | Median $c$ | Median $s$ | $d_q(i)$: median (IQR) |
| --- | --- | --- | --- | --- | --- |
| $R_1$ | Coala | 100.0 | 0.800 | 0.020 | 0.000 (0.000) |
|  | treeducken | 7.9 | 0.430 | 0.000 | 0.070 (0.063) |
|  | cophylo | 100.0 | 0.800 | 0.025 | 0.000 (0.000) |
|  | AsymmeTree | 71.3 | 0.552 | 0.012 | 0.000 (0.014) |
| $R_2$ | Coala | 100.0 | 0.080 | 0.580 | 0.000 (0.000) |
|  | treeducken | 36.5 | 0.000 | 0.366 | 0.034 (0.133) |
|  | cophylo | 100.0 | 0.076 | 0.601 | 0.000 (0.000) |
|  | AsymmeTree | 76.7 | 0.191 | 0.556 | 0.000 (0.000) |
| $R_3$ | Coala | 89.5 | 0.306 | 0.306 | 0.000 (0.000) |
|  | treeducken | 57.3 | 0.237 | 0.222 | 0.000 (0.016) |
|  | cophylo | 100.0 | 0.301 | 0.298 | 0.000 (0.000) |
|  | AsymmeTree | 60.2 | 0.346 | 0.313 | 0.000 (0.026) |

**Figure 2.**
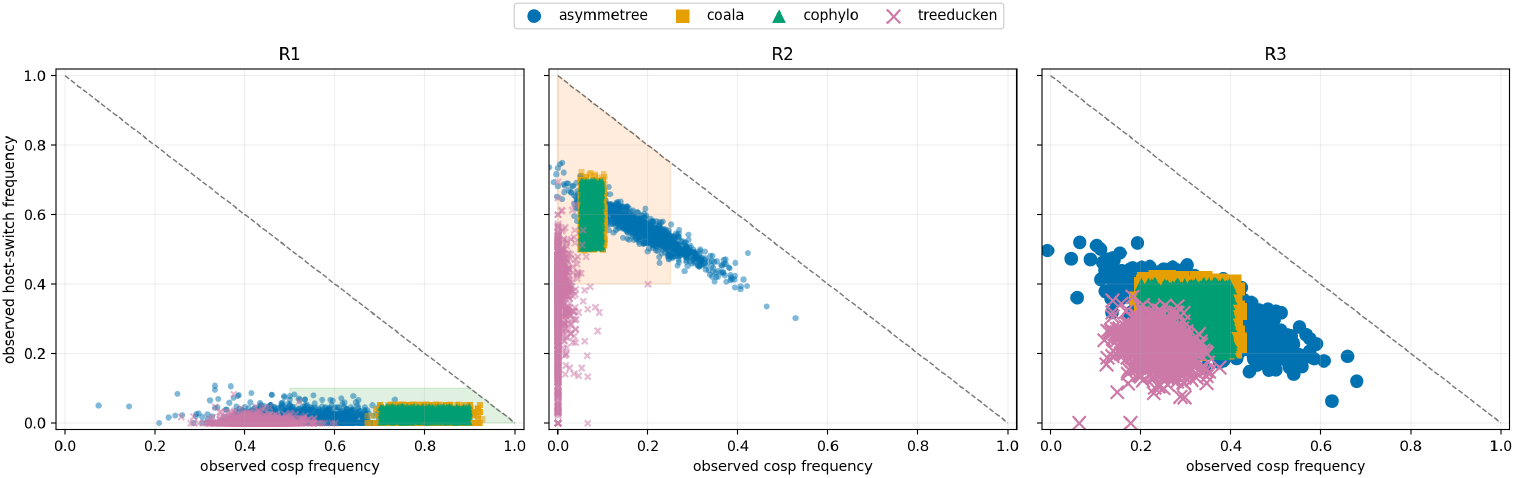
Observed cospeciation and host-switch frequencies for the datasets generated under the three simulation regimes (as columns). Each point represents one simulated dataset, and colours distinguish the four generators. The shaded areas correspond to the intended regions *R*_1_, *R*_2_, and *R*_3_.

**Figure 3.**
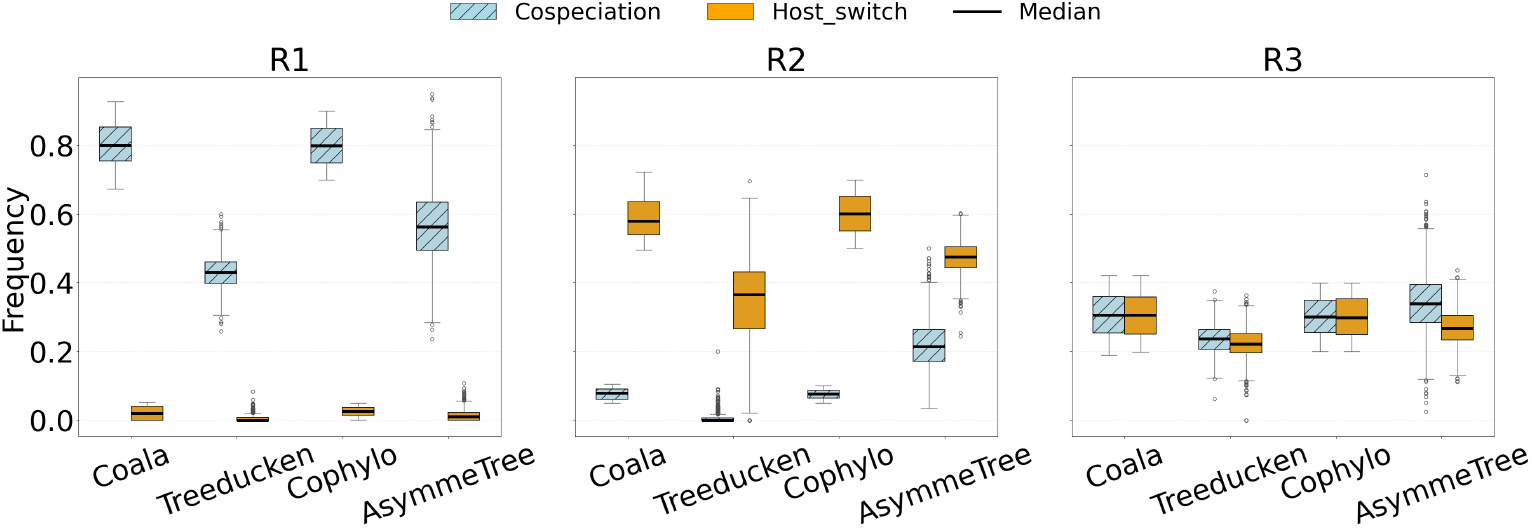
Boxplots of observed cospeciation and host-switch frequencies across generators in regimes *R*_1_-*R*_3_.

**Figure 4.**
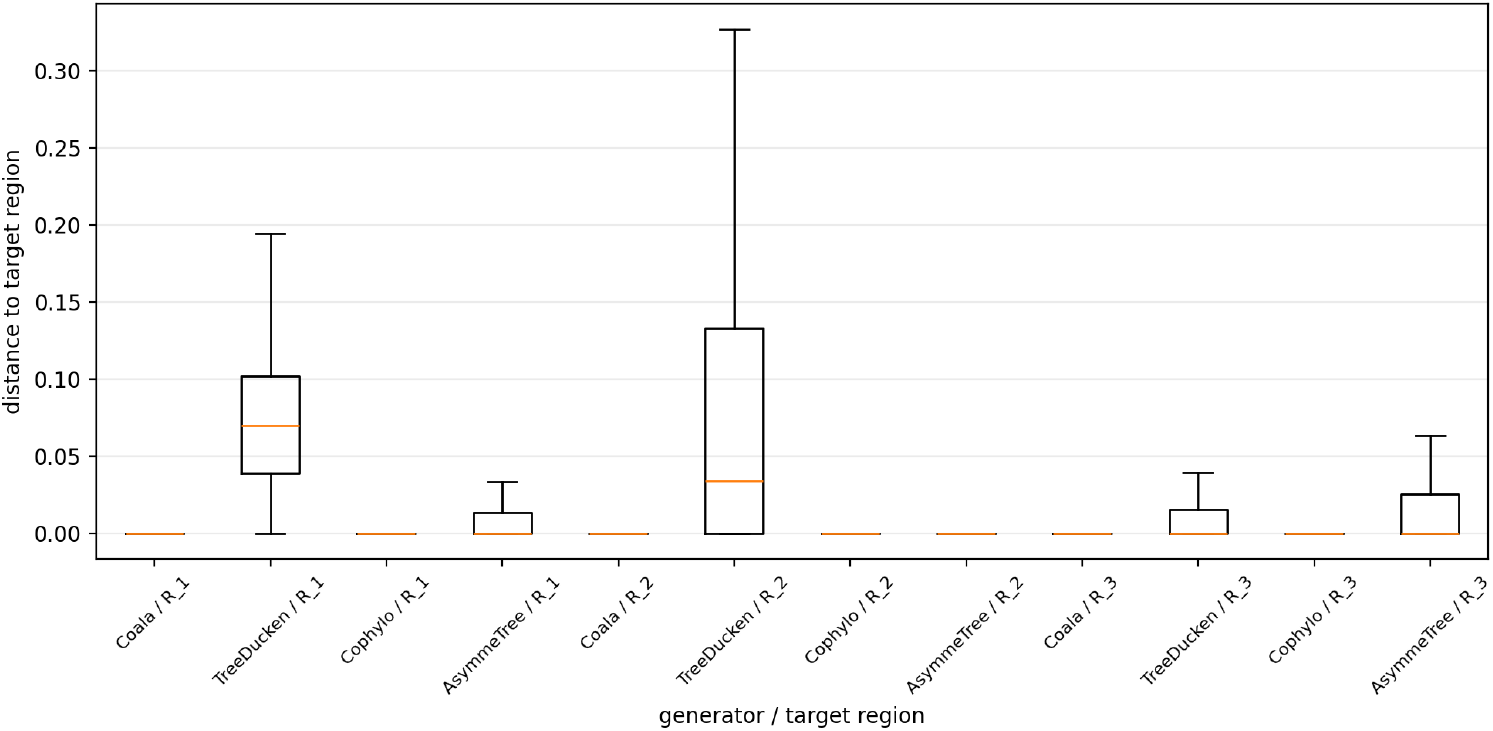
Distribution of the Euclidean distance 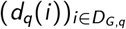 between each simulated dataset *D*_*G,q*_ and the region corresponding to its intended regime *R*_*q*_, shown for each generator and regime.

In the cospeciation-dominated regime *R*_1_, all datasets generated by `Coala` and `cophylo` fall inside the target region. `AsymmeTree` also performs well, with 71.3% of its datasets inside *R*_1_. The median observed pairs (*c*_*i*_, *s*_*i*_) are (0.800, 0.020) for `Coala`, (0.800, 0.025) for `cophylo`, and (0.552, 0.012) for `AsymmeTree`. The median distances from *R*_1_ are all zero for these three generators, with zero IQR for both `Coala` and `cophylo` and a small 0.014 IQR for `AsymmeTree`. In contrast, `treeducken` also produces very low host-switch frequencies in *R*_1_, but its cospeciation frequency is usually below the threshold *c* = 0.50. Its median observed (*c, s*) pair is (0.430, 0), and only 7.9% of its datasets fall inside *R*_1_. The median distance from the region is 0.070, with an IQR of 0.063. This result is linked to the way `treeducken` parameters have been set (see section 2.2). With the parameter settings used here, increasing the cospeciation rate also increases the host and symbiont extinction rates. The observed cospeciation frequency therefore does not increase at the same rate. In preliminary simulations, fixing the extinction rates independently while increasing the cospeciation rate resulted in trees with extremely large numbers of leaves, making the simulations impractical. Thus, in `treeducken`, obtaining very high cospeciation frequencies while keeping the trees computationally manageable is difficult.

In the host-switch-dominated regime *R*_2_, all datasets generated by `Coala` and `cophylo` fall inside the target region. Their median observed pairs (*c*_*i*_, *s*_*i*_) are (0.080, 0.580) and (0.076, 0.601), respectively, and their median distances from *R*_2_ are zero (with zero IQR). `AsymmeTree` falls inside *R*_2_ in 76, 7% of the simulations. Its median observed pair is (0.191, 0.556). Thus the host-switch frequency is high, while the cospeciation frequency is only slightly below the upper boundary *c* = 0.25. Its median distance from *R*_2_ is also zero (with zero IQR). Nevertheless, only 76.7% of the simulated datasets fall within the target region, reflecting a wider dispersion around the target than for `Coala` and `cophylo`.

Contrasting again with these results, `treeducken` has only 36.5% of the datasets falling inside *R*_2_. Its median observed pair (*c*_*i*_, *s*_*i*_) = (0, 0.366) lies outside all three regimes. In particular, although cospeciation remains low, the host-switch frequency is below the threshold *s* = 0.40 required for *R*_2_. The median distance from *R*_2_ is 0.034, with an IQR of 0.133.

In the mixed regime *R*_3_, `cophylo` succeeds in having all of its datasets inside the target region. It is followed by `Coala` with 89.5% of attainment proportion. Their median observed pairs (*c*_*i*_, *s*_*i*_) are (0.301, 0.298) for `cophylo` and (0.306, 0.306) for `Coala`. Their median distances from *R*_3_ and IQR are all zero. `AsymmeTree` has only 60.2% of its datasets falling inside *R*_3_. Its median observed pair is (0.346, 0.313), with the cospeciation frequency slightly below the upper boundary *c* = 0.40. Its median distance from *R*_3_ is zero with a small IQR of 0.026, showing that some datasets lie close to, but outside, the region. `treeducken` has the lowest proportion of datasets inside *R*_3_ equal to 57.3%. Its median observed frequencies pair, (0.237, 0.222), lies inside the region, and its median distance is zero, with small IQR 0.016.

### 3.2. Generators comparison

#### 3.2.1. Tree-based analysis

Figs. 5 and 6 show how strongly the regime parameters overall constrain tree growth across the different regimes and generators.

**Figure 5.**
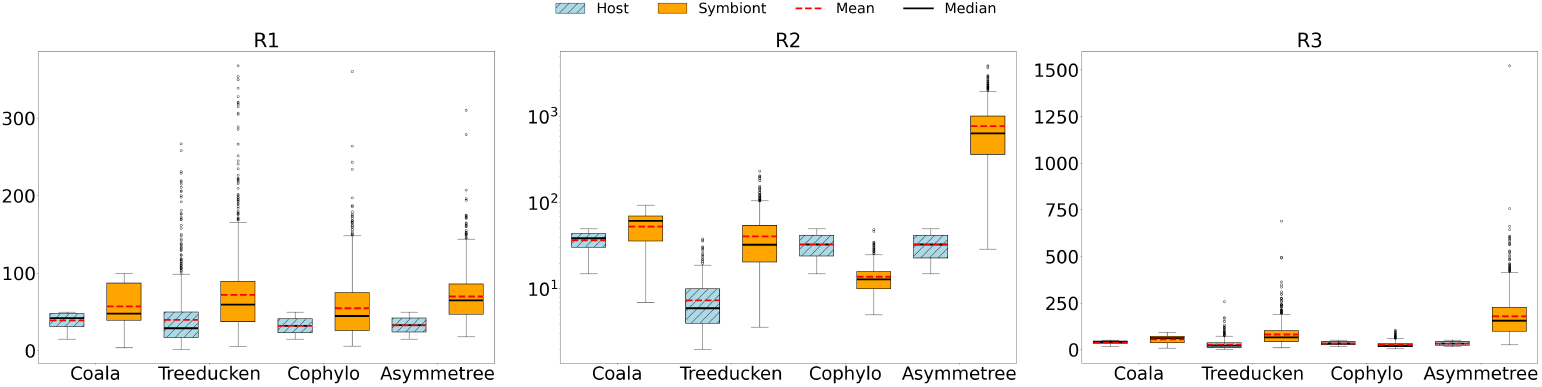
Boxplots of the number of leaves for each regime *R*_1_-*R*_3_

**Figure 6.**
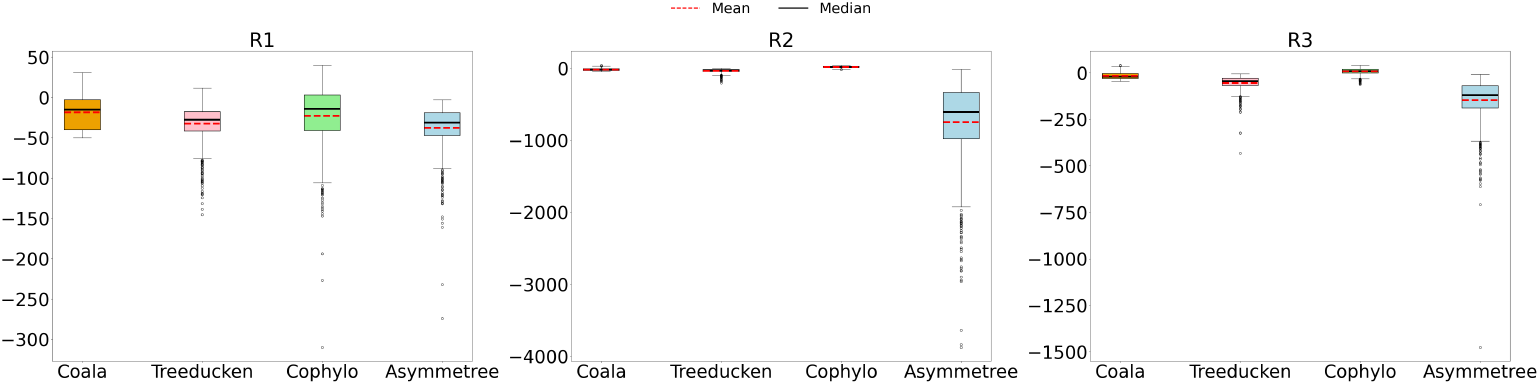
Boxplots of the difference in the number of leaves between host and symbiont trees for each simulated dataset, across regimes *R*_1_-*R*_3_.

Let us recall that for `Coala`, `cophylo` and `AsymmeTree` generators, host trees were generated as a first step, with sizes that could thus be controlled, while `treeducken` had host trees jointly generated with the symbiont trees. This is particularly striking in regime *R*_1_, where host trees for `treeducken` can be very large, due to the high co-speciation values. In regime *R*_1_, high cospeciation and low host-switch should keep symbiont and host tree sizes similar, and this is indeed what we observe across all four generators.

In *R*_2_, frequent switches allow symbionts to diversify independently, and this may lead to a dramatic increase in symbiont size for `AsymmeTree` and to a lesser extent for `treeducken`, while `Coala` and `cophylo`’s symbiont trees remain constrained.

In *R*_3_, differences are smaller but `AsymmeTree` again, generates the largest trees, showing that its growth is less tightly controlled by the regime.

Finally, especially in the *R*_2_ and *R*_3_ regimes, looking at the difference in the number of leaves between host and symbiont, we observe that the simulated symbiont trees are often larger than the corresponding host trees, indicating that under these parameter settings, symbiont lineages diversify more rapidly than host lineages. In *R*_1_, this effect is less pronounced overall, but it is still very visible for `AsymmeTree`, where symbiont trees can grow substantially even under high cospeciation. This behaviour is expected in that model, which was originally designed for gene-species evolution, where gene lineages can diversify more independently from the species tree.

The three indices from Figure 7 capture different aspects of tree balance. A higher Cherry index means that the tree contains more pairs of sibling leaves, which is generally associated with greater local balance; conversely, a lower Cherry index indicates fewer such pairs and is associated with greater imbalance. The Sackin index summarizes the total root-to-leaf depth of the tree; for a fixed number of leaves, lower Sackin values are associated with more balanced trees, whereas higher values indicate greater imbalance. Lower Colless values indicate that internal splits tend to divide the tree into subtrees of similar size, while higher Colless values indicate more asymmetric splits and therefore greater imbalance. Thus, higher Cherry together with lower Sackin and Colless values provides a consistent indication of a more balanced tree, whereas lower Cherry together with higher Sackin and Colless values indicates a more imbalanced tree.

**Figure 7.**
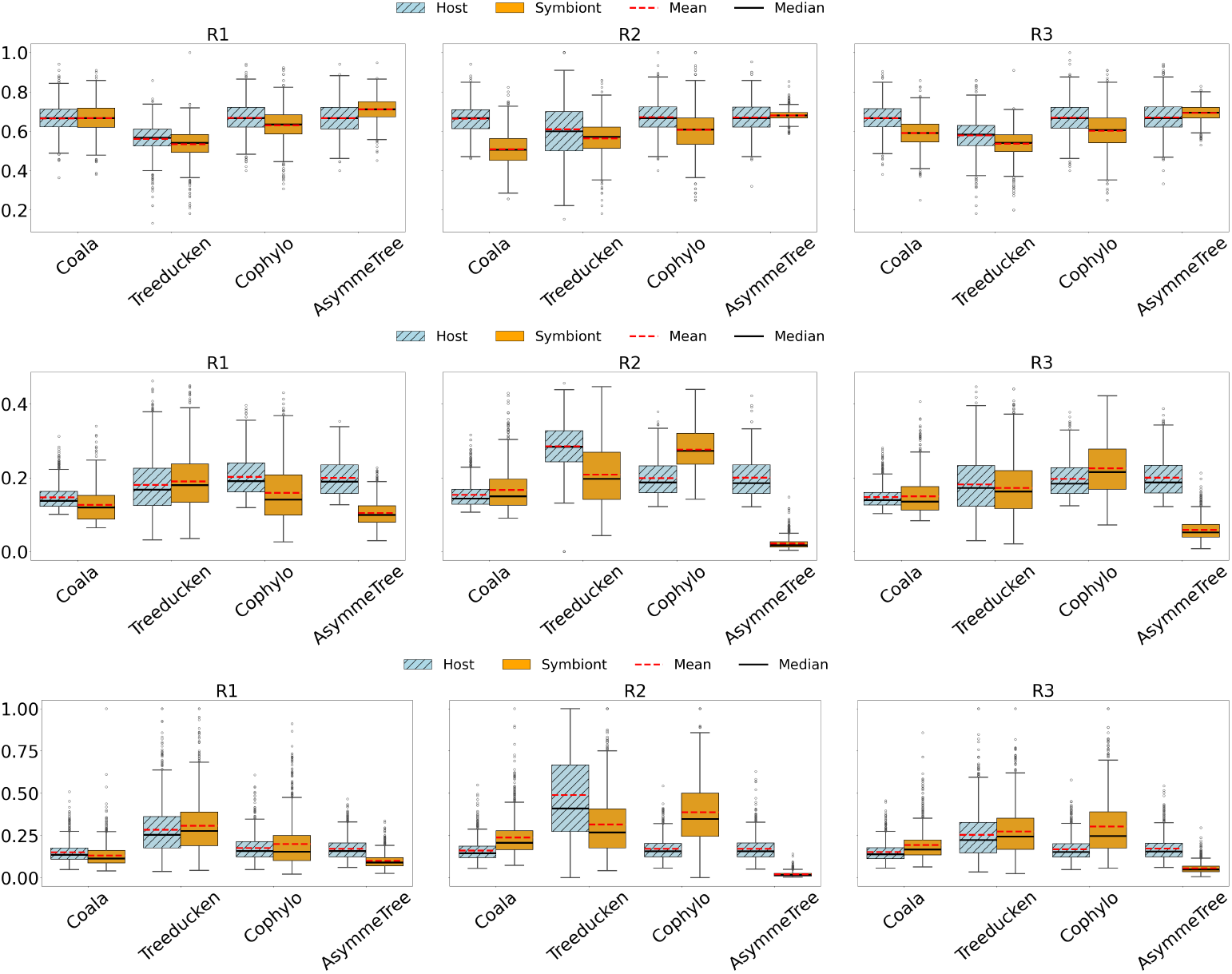
Normalized Cherry (first row), Sackin (second row) and Colless (third row) indices across regimes *R*_1_-*R*_3_ (in columns).

In the high-cospeciation regime (*R*_1_), host and symbiont tree-shape indices are generally similar for `Coala` and `treeducken`. For `cophylo`, the indices provide a mixed picture: the symbiont tree has lower Sackin and Colless values, but also a lower Cherry index. In contrast, `AsymmeTree` shows a consistent shift towards more balanced symbiont trees, with a higher Cherry index and substantially lower Sackin and Colless indices.

In the switch-dominated regime (*R*_2_), the effect on tree shape becomes strongly generator-dependent. `Coala` and especially `cophylo` show lower Cherry and higher Sackin and Colless values for the symbiont tree, consistently indicating greater imbalance. `treeducken` shows a mixed pattern, since the Cherry index decreases while Sackin and Colless also decrease. In contrast, `AsymmeTree` exhibits substantially lower Sackin and Colless values for the symbiont tree, indicating a markedly different tree-shape pattern.

The mixed regime (*R*_3_) shows a pattern similar to *R*_2_, although generally less pronounced. `Coala` and `cophylo` tend to produce symbiont trees with lower Cherry and higher Sackin and Colless values, whereas `treeducken` again shows a mixed signal across the three indices. `AsymmeTree` retains the opposite pattern, with similar or slightly higher Cherry values and substantially lower Sackin and Colless indices for the symbiont tree.

Thus, the effect of the coevolutionary regime on tree shape is strongly generator-dependent. In particular, `cophylo` and `AsymmeTree` often display opposite host-symbiont shape patterns in *R*_2_ and *R*_3_, while `treeducken` frequently shows different signals across the three balance indices.

#### 3.2.2. Association-based measures

In Table 6 and Figs. 8-9, the network-level measures show that each generator produces markedly different host-symbiont association structures across regimes, reflecting how associations are constructed in the simulation. In `Coala`, each symbiont is associated with exactly one host, which keeps the association network sparse across regimes.

**Table 6.** Mean, Median, and Standard Deviation (sd) of association network density across regimes *R*_1_-*R*_3_.

|  | Mean/Median (sd) | Mean/Median (sd) | Mean/Median (sd) |
| --- | --- | --- | --- |
| Method | $R_1$ | $R_2$ | $R_3$ |
| Coala | 0.033 / 0.029 (0.014) | 0.041 / 0.032 (0.019) | 0.041 / 0.033 (0.019) |
| treeducken | 0.140 / 0.091 (0.145) | 0.416 / 0.343 (0.251) | 0.164 / 0.111 (0.169) |
| cophylo | 0.117 / 0.077 (0.123) | 0.240 / 0.229 (0.085) | 0.158 / 0.132 (0.097) |
| AsymmeTree | 0.034 / 0.030 (0.012) | 0.034 / 0.030 (0.013) | 0.034 / 0.030 (0.013) |

**Figure 8.**
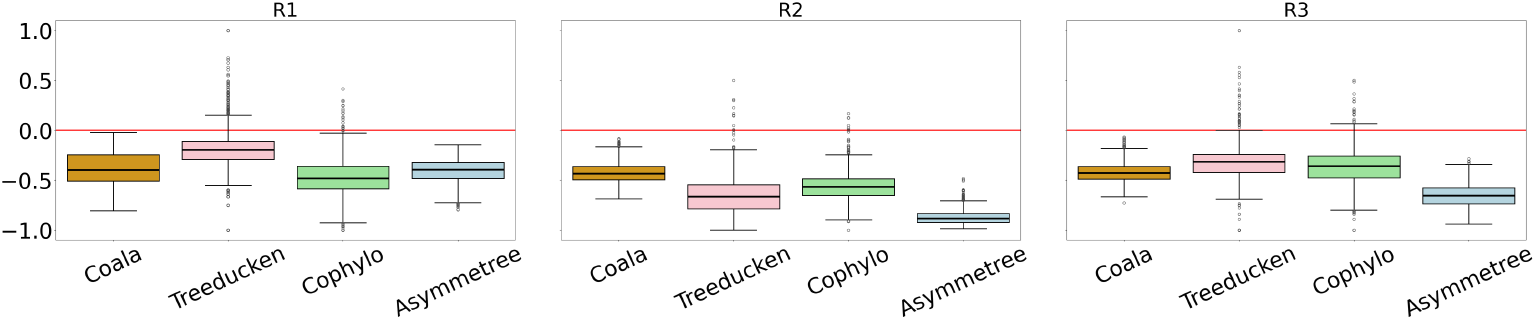
Degree assortativity across regimes *R*_1_-*R*_3_.

**Figure 9.**
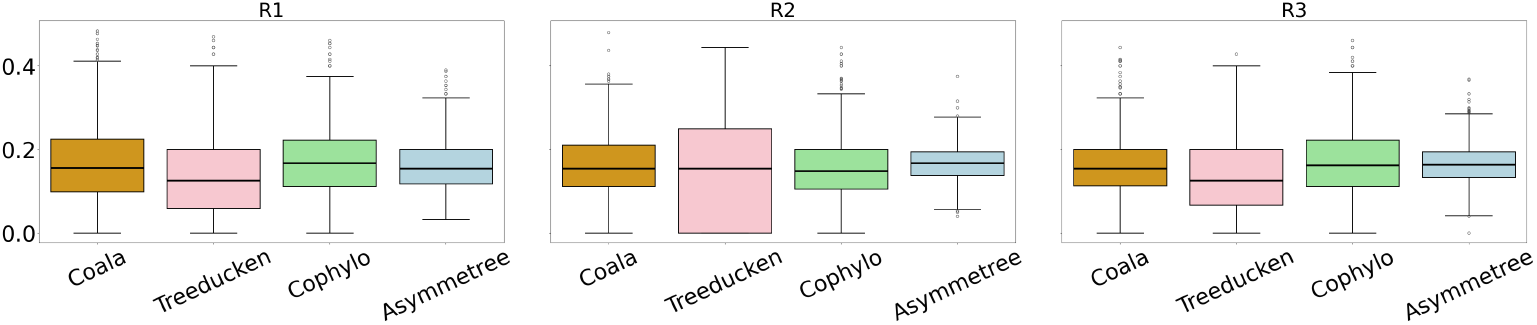
Host hotspots frequency across regimes *R*_1_-*R*_3_.

Thus, the degree variation comes mainly from hosts accumulating symbionts, so the proportion of host hotspots remains similar in *R*_1_-*R*_3_. Degree assortativity is consistently negative, indicating that edges tend to link nodes with different degrees. Because each symbiont can attach to only one host, this dis-assortative structure is expected and remains stable across regimes. In `treeducken`, density is low under high cospeciation and in the mixed regime (medians of 0.091 and 0.111 respectively) but roughly three times higher when switching dominates (0.343). Degree assortativity is negative everywhere and tracks the switching rate (median values of −0.196, −0.666 and −0.316 in *R*_1_-*R*_3_): frequent switching concentrates associations on a few high-degree hosts, so edges link nodes with increasingly different degrees. Host hotspots, by contrast, stay essentially constant (median values of 0.125, 0.148, 0.161 in *R*_1_-*R*_3_), suggesting they reflect the generator rather than the regime. In `cophylo`, density behaves similarly to `treeducken` by staying lower under high-cospeciation and mixed regime but twice higher when moving to high-switch regime *R*_2_. Degree assortativity is low in the three regimes and hosts hotspots distribution is stable across them. In `AsymmeTree`, regime does not affect the density distribution, which stays low everywhere. Degree assortativity is negative in all regimes, and it is most negative in *R*_2_. This matches the regime meaning: when switching dominates, links are more likely to connect nodes with very different degrees, which pushes assortativity down; when cospeciation dominates, this effect is weaker and assortativity moves closer to zero. Thus, the distribution of host hotspots shows little variation across regimes.

### 3.3. Effect of the generator on parsimonious reconciliation

In *Q*_2_, we asked whether differences between generators also appear in the output of a standard downstream analysis. We therefore compared the optimal parsimonious event profiles obtained with the same cost vector within each regime.

#### 3.3.1. Optimal event profiles

Figure 10 gives a dataset-level view of the optimal parsimonious event profiles. Each dataset is represented by the mean of the profiles associated with its distinct optimal event vectors. Table 7 reports the corresponding means across datasets.

**Table 7.** Mean dataset-level optimal event profiles. Values are the mean cospeciation and host-switch proportions 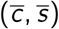 across datasets for each generator, regime, and cost vector.

| Regime | Cost vector | Coala | treeducken | AsymmeTree | cophylo |
| --- | --- | --- | --- | --- | --- |
| R1 | $\langle 0, 1, 2, 1 \rangle$ | (0.8458, 0.0263) | (0.7408, 0.0716) | (0.6345, 0.0355) | (0.4779, 0.1389) |
| | $\langle 0, 1, 3, 1 \rangle$ | (0.8417, 0.0216) | (0.7171, 0.0372) | (0.6210, 0.0147) | (0.4593, 0.0862) |
| R2 | $\langle 0, 1, 2, 1 \rangle$ | (0.2199, 0.6082) | (0.2331, 0.2267) | (0.3508, 0.4033) | (0.3669, 0.3086) |
| | $\langle 0, 1, 3, 1 \rangle$ | (0.2332, 0.4883) | (0.2256, 0.1101) | (0.3390, 0.3457) | (0.3543, 0.2207) |
| R3 | $\langle 0, 1, 2, 1 \rangle$ | (0.3924, 0.3685) | (0.4550, 0.2865) | (0.4506, 0.2494) | (0.3760, 0.2790) |
| | $\langle 0, 1, 3, 1 \rangle$ | (0.3810, 0.2915) | (0.4348, 0.2103) | (0.4388, 0.2142) | (0.3601, 0.1921) |

**Figure 10.**
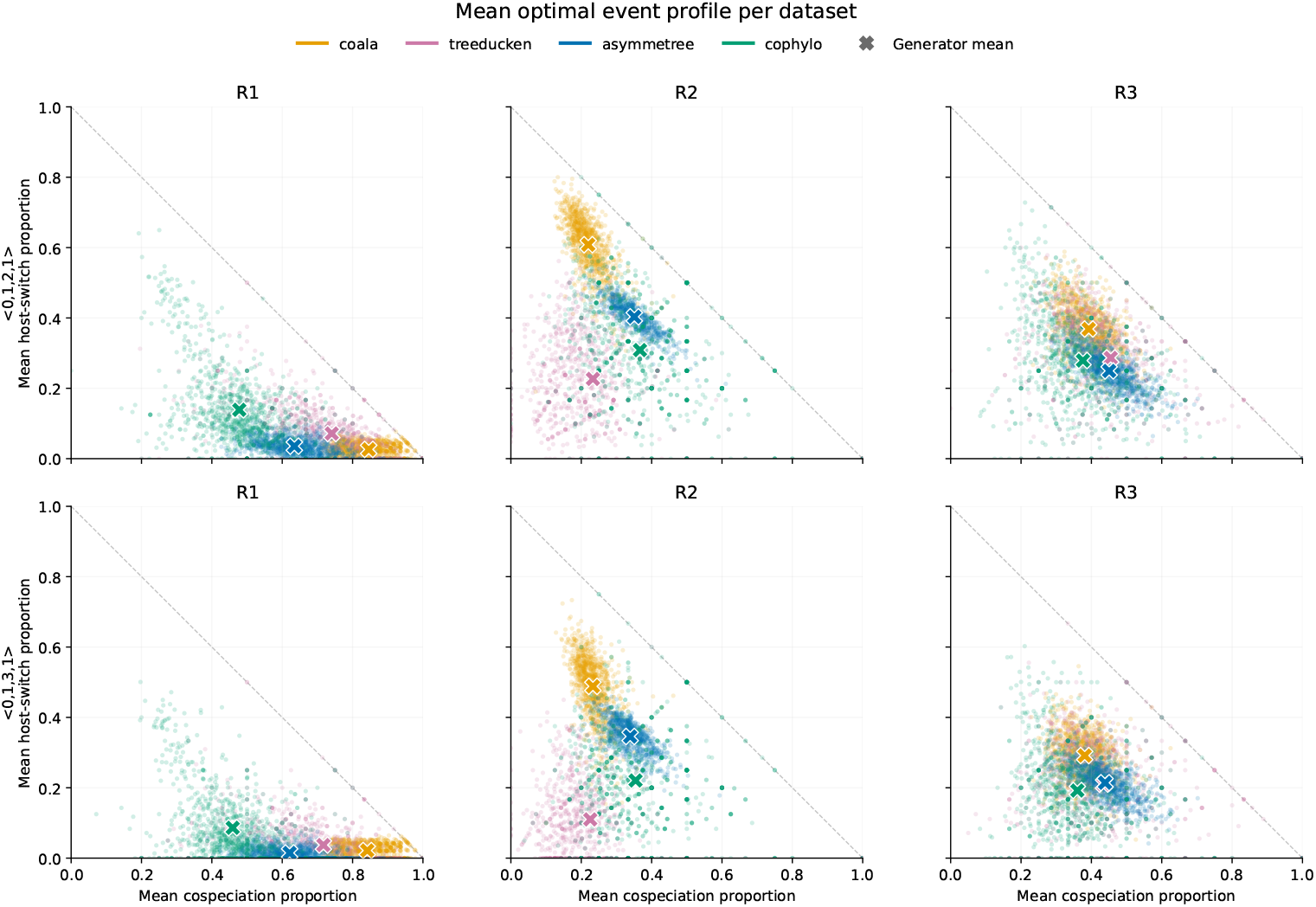
Mean optimal parsimonious event profile for each dataset. Each point represents one dataset and is obtained by averaging the profiles associated with its distinct optimal event vectors. Crosses show the mean of these dataset-level profiles for each generator. Columns correspond to regimes R1–R3 and rows to the two reconciliation cost vectors.

In *R*_1_, the profiles are generally concentrated towards high-cospeciation and low host-switch proportions, although their patterns differ across generators. `Coala` is the most concentrated towards high cospeciation. `AsymmeTree` is also concentrated at low host-switch values, but at lower cospeciation proportions than `Coala`. As for `treeducken`, it has a broader distribution and includes more profiles with larger host-switch proportions, while `cophylo`shows an even broader spread towards lower cospeciation and higher host-switch proportions.

The differences are strongest in *R*_2_. With the cost vector ⟨0, 1, 2, 1⟩, `Coala` produces profiles with high host-switch proportions, while `AsymmeTree` is concentrated around more comparable cospeciation and host-switch proportions. `treeducken` shows a different pattern, with substantially lower host-switch proportions. `cophylo` occupies an intermediate region, with substantial overlap with `AsymmeTree` but generally lower host-switch proportions than `Coala`. These differences remain when the host-switch cost is increased from 2 to 3.

In *R*_3_, the dataset-level mean profiles overlap more strongly across generators. However, `Coala` still tends towards larger host-switch proportions, while `AsymmeTree` tends towards larger cospeciation and lower host-switch proportions. `treeducken` and `cophylo` occupy largely overlapping intermediate regions.

The results show that datasets produced under the same intended regime can therefore lead to different optimal event compositions when analysed with the same parsimonious reconciliation method.

The generator-level means reported in Table 7 confirm these differences. They are most pronounced in *R*_2_: with cost vector ⟨0, 1, 2, 1⟩, the mean profiles 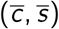 are (0.2199, 0.6082) for `Coala`, (0.2331, 0.2267) for `treeducken`, (0.3508, 0.4033) for `AsymmeTree`, and (0.3669, 0.3086) for `cophylo`. Increasing the host-switch cost to 3 reduces the host-switch proportion for all four generators, while the differences among generators remain.

#### 3.3.2. Number of optimal event profiles

The generators also differ in the number of distinct optimal event profiles compatible with the minimum parsimony cost (Figure 11 and Table 8).

**Table 8.** Ambiguity of the optimal parsimonious event profiles. For each generator, we report the median number of distinct optimal event profiles (*c, s*), with the interquartile range in square brackets.

| Regime | Cost vector | Coala | treeducken | AsymmeTree | cophylo |
| --- | --- | --- | --- | --- | --- |
| R1 | $\langle 0, 1, 2, 1 \rangle$ | 1 [1, 2] | 1 [1, 2] | 3 [2, 5] | 3 [2, 5] |
| | $\langle 0, 1, 3, 1 \rangle$ | 1 [1, 1] | 1 [1, 2] | 1 [1, 2] | 1 [1, 2] |
| R2 | $\langle 0, 1, 2, 1 \rangle$ | 6 [3, 11] | 2 [1, 4] | 124 [52, 244] | 1 [1, 2] |
| | $\langle 0, 1, 3, 1 \rangle$ | 4 [2, 7] | 1 [1, 2] | 78 [28, 173] | 1 [1, 2] |
| R3 | $\langle 0, 1, 2, 1 \rangle$ | 6 [2, 11] | 2 [1, 4] | 10 [4, 24] | 2 [1, 3] |
| | $\langle 0, 1, 3, 1 \rangle$ | 3 [2, 4] | 1 [1, 2] | 4 [2, 8] | 1 [1, 2] |

**Figure 11.**
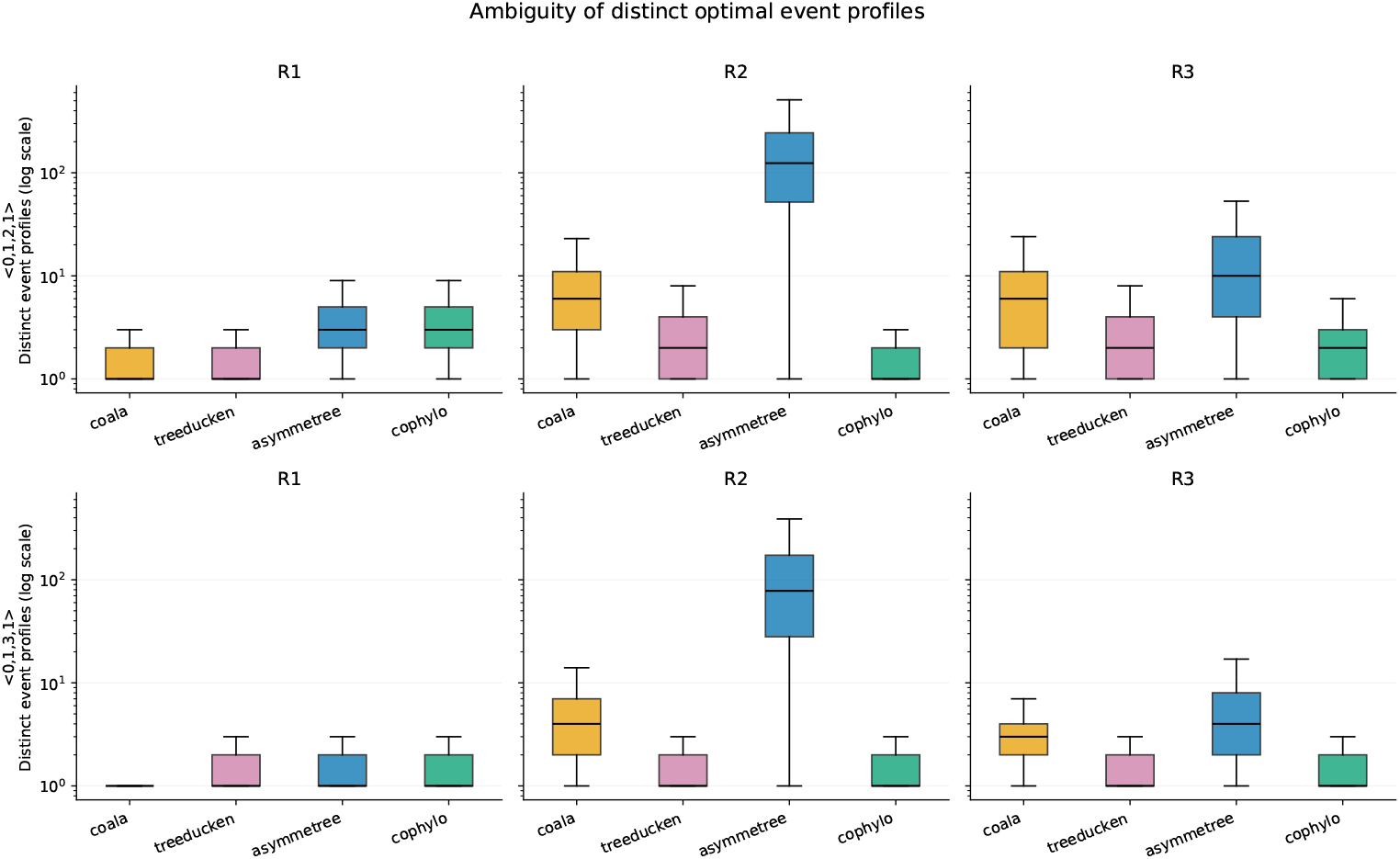
Number of distinct optimal event profiles for each generator, regime, and cost vector. The *y* axis is shown on a logarithmic scale. Columns correspond to regimes *R*_1_-*R*_3_ and rows to the two reconciliation cost vectors.

In *R*_1_, ambiguity at the event-profile level is generally limited. With cost vector ⟨0, 1, 2, 1⟩, the median number of distinct optimal event profiles is 1 for `Coala` and `treeducken`, and 3 for `AsymmeTree` and `cophylo`. With ⟨0, 1, 3, 1⟩, the median is 1 for all four generators.

The largest differences occur in *R*_2_. With cost vector ⟨0, 1, 2, 1⟩, the median number of distinct optimal event profiles is 6 for `Coala`, 2 for `treeducken`, 124 for `AsymmeTree`, and 1 for `cophylo`. When the host-switch cost is increased to 3, the corresponding values are 4, 1, 78, and 1, respectively.

*R*_3_ gives intermediate values. With cost vector ⟨0, 1, 2, 1⟩, the medians are 6, 2, 10, and 2 for `Coala`, `treeducken`, `AsymmeTree`, and `cophylo`, respectively. With ⟨0, 1, 3, 1⟩, they decrease to 3, 1, 4, and 1.

#### 3.3.3. Datasets included in the analysis

Table 9 reports the number of datasets for which the downstream reconciliation analysis completed within the imposed running-time limit. Most datasets were successfully analysed, although the completion rate varied across generators, regimes, and cost vectors.

**Table 9.** Number of datasets used in the downstream reconciliation analysis.

| Regime | Cost vector | Coala | treeducken | cophylo | AsymmeTree |
| --- | --- | --- | --- | --- | --- |
| R1 | $\langle 0, 1, 2, 1 \rangle$ | 1000 | 989 | 989 | 1000 |
| | $\langle 0, 1, 3, 1 \rangle$ | 1000 | 990 | 989 | 1000 |
| R2 | $\langle 0, 1, 2, 1 \rangle$ | 1000 | 854 | 990 | 697 |
| | $\langle 0, 1, 3, 1 \rangle$ | 1000 | 854 | 990 | 844 |
| R3 | $\langle 0, 1, 2, 1 \rangle$ | 1000 | 977 | 994 | 999 |
| | $\langle 0, 1, 3, 1 \rangle$ | 1000 | 977 | 994 | 1000 |

The largest reduction occurs in *R*_2_, particularly for `AsymmeTree`: 697 datasets completed with cost vector ⟨0, 1, 2, 1⟩ and 844 with ⟨0, 1, 3, 1⟩. For `treeducken`, 854 datasets completed under both cost vectors, while `cophylo` retained 990 and `Coala` all 1000 datasets. The lower completion rates are associated with instances having very large optimal-solution spaces, for which the reconciliation computation exceeded the imposed running-time limit.

Consequently, the ambiguity summaries reported above are computed only on successfully completed runs and should be interpreted with this computational filtering in mind.

Taken together, these results answer *Q*_2_ positively. Differences among generators are also visible in a standard downstream parsimonious reconciliation analysis. They affect both the inferred event composition and, in some regimes, the number of equally optimal solutions. The effect is particularly strong in the switch-dominated regime *R*_2_ and remains visible with both reconciliation cost vectors.

### 3.4. Comparison to real data

The comparison included a reference set of 30 real datasets summarized through the eight tree-based features and 1000 synthetic datasets for each generator-regime combination. Figure 12 shows the projection obtained by fitting PCA on the real datasets and projecting the synthetic datasets onto the same first two axes. Those first two components explain 79.1% of the variation in the real datasets.

**Figure 12.**
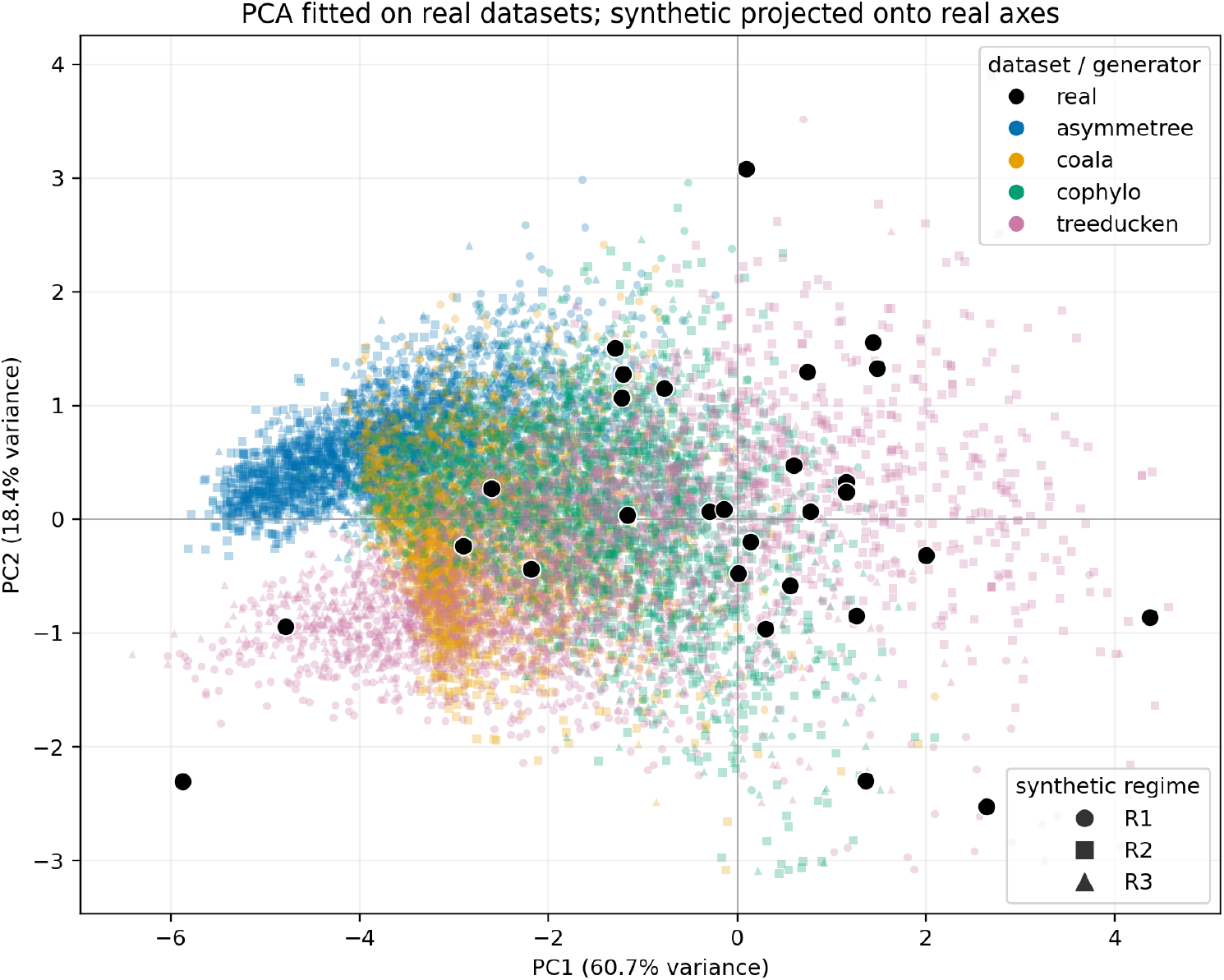
Projection of the real and synthetic datasets onto the first two principal components. The PCA was fitted on the standardized real datasets, and the synthetic datasets were projected onto the same axes. Colours identify the generators and symbols identify the simulation regimes. PC1 and PC2 explain 60.7% and 18.4% of the variation in the real datasets, respectively.

The positions of the synthetic datasets depend strongly on the generator. `Coala` produces a relatively compact set of points, while `cophylo` and `treeducken` cover a wider area. `AsymmeTree` changes markedly across regimes. In particular, many of its datasets from *R*_2_ lie far from the real datasets. The PCA is used only as a visualization. The following quantitative comparison is based on distances computed in the full eight-dimensional feature space.

#### 3.4.1. Proximity of synthetic datasets to real data

Table 10 reports, for each generator and regime, the distance from each synthetic dataset to its closest real dataset, together with the corresponding proximity rate.

**Table 10.**
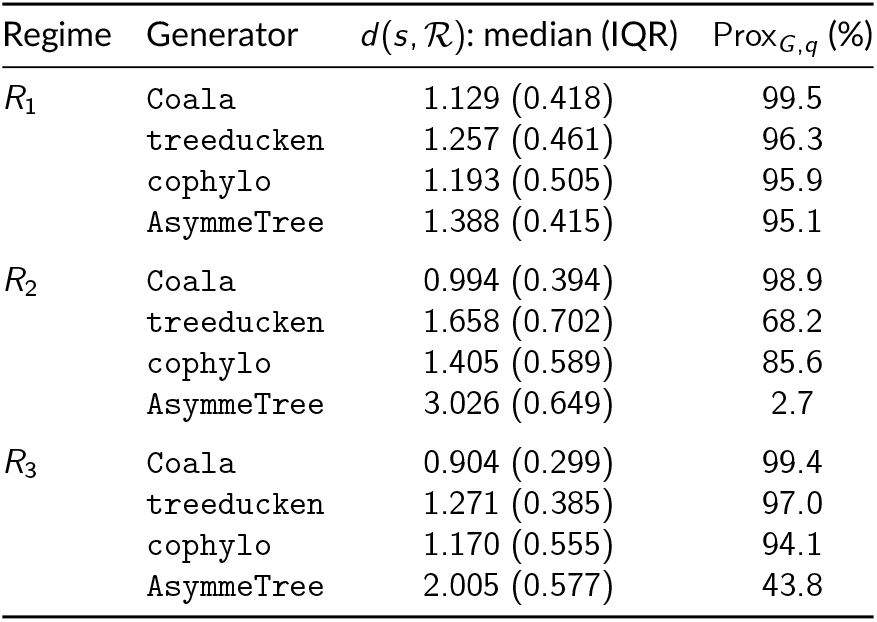
Proximity of synthetic datasets to the real reference set ℛ. For each generator and regime, we report the median distance from a synthetic dataset *s* to its closest real dataset, *d* (*s*, ℛ), with the interquartile range (IQR) in parentheses. We also report the proximity rate Prox_*G,q*_, defined as the percentage of synthetic datasets satisfying *d* (*s*, ℛ) ≤ *τ*, with *τ* = 1.945. Lower distances and higher proximity rates indicate greater similarity to the real reference set.

`Coala` has the lowest median distance in all three regimes, equal to 1.129 in *R*_1_, 0.994 in *R*_2_, and 0.904 in *R*_3_. Its proximity rate is always above 98%, indicating that almost all datasets generated by `Coala` are close to at least one dataset in the real reference set.

`treeducken` also shows high proximity in *R*_1_ and *R*_3_, with proximity rates of 96.3% and 97.0%, respectively. In *R*_2_, however, its median distance increases to 1.658 and its proximity rate decreases to 68.2%.

`cophylo` shows high proximity to the real reference set in all three regimes. Its proximity rates are 95.9% in *R*_1_, 85.6% in *R*_2_, and 94.1% in *R*_3_. Although its median distances are generally higher than those of `Coala`, the large majority of its simulated datasets remain within the empirical proximity threshold.

`AsymmeTree` shows the strongest dependence on the simulation regime. In *R*_1_, its proximity rate is 95.1%, whereas it decreases to 43.8% in *R*_3_ and to only 2.7% in *R*_2_. Consistently, the median distance in *R*_2_ is 3.026, the largest among all generator-regime combinations.

#### 3.4.2. Coverage of the real datasets

Table 11 reports, for each generator and regime, the distance from each real dataset to its closest synthetic dataset, together with the corresponding coverage rate.

**Table 11.**
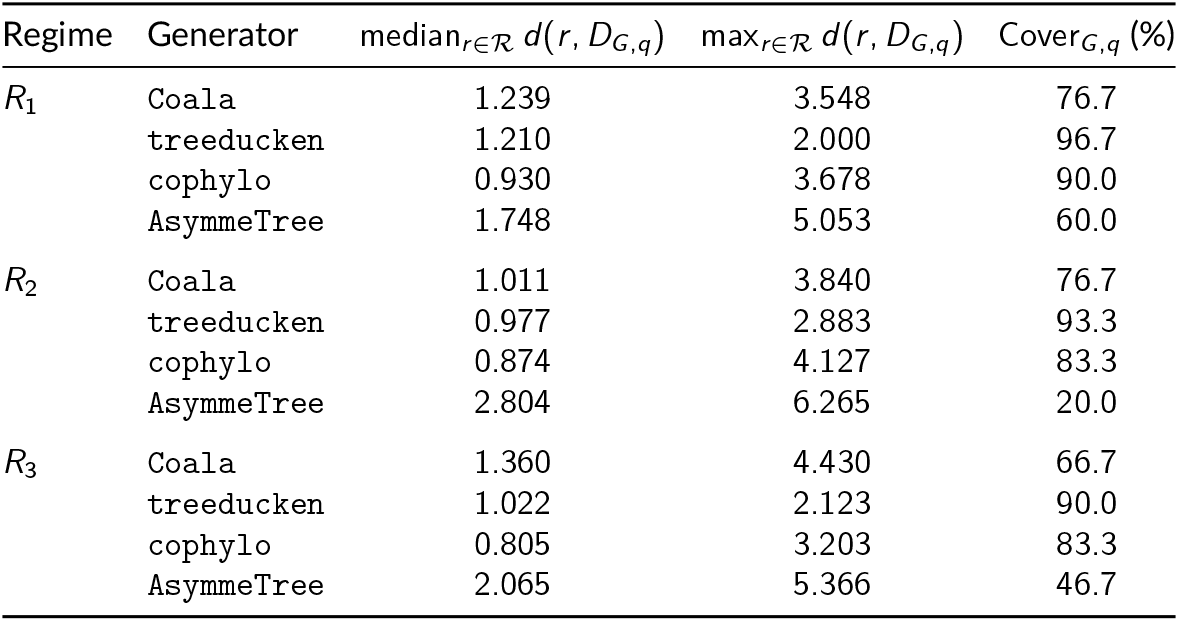
Coverage of the reference set ℛ. The table reports the median and maximum of the distances *d* (*r, D*_*G,q*_) of any reference dataset *r* to the closest synthetic dataset from the corresponding generator and regime *D*_*G,q*_, as well as the coverage rate Cover_*G,q*_. The coverage rate is the percentage of real datasets with distance at most *τ* = 1.945. Lower distances and higher percentages indicate broader coverage of the reference set ℛ.

`treeducken` provides the highest coverage in all three regimes, with coverage rates of 96.7% in *R*_1_, 93.3% in *R*_2_, and 90.0% in *R*_3_. It also has the lowest maximum distance in every regime, indicating that almost all real datasets have a relatively close synthetic counterpart generated by `treeducken`.

`cophylo` also provides broad coverage of the reference set, with coverage rates of 90.0% in *R*_1_ and 83.3% in both *R*_2_ and *R*_3_. Moreover, it has the lowest median real-to-synthetic distance in all three regimes, equal to 0.930, 0.874, and 0.805, respectively. Thus, while `treeducken` covers the largest fraction of the reference set, `cophylo` tends to provide particularly close synthetic counterparts for the real datasets that it represents.

`Coala` shows lower coverage, ranging from 66.7% in *R*_3_ to 76.7% in *R*_1_ and *R*_2_. This contrasts with its very high proximity rates: almost all datasets generated by `Coala` are close to at least one real dataset, but they cover a more restricted portion of the reference set.

`AsymmeTree` again shows the strongest dependence on the regime. Its coverage is 60.0% in *R*_1_ and 46.7% in *R*_3_, and decreases to only 20.0% in *R*_2_. In *R*_2_, it also has the largest median and maximum distances, equal to 2.804 and 6.265, respectively.

#### 3.4.3. Differences between individual features

Figure 13 shows the median standardized value of each feature for every generator and regime. Each feature is standardized using the mean and standard deviation computed across the 30 real datasets. Thus, a value of zero corresponds to the empirical mean, while values of +1 and −1 correspond to one empirical standard deviation above and below that mean, respectively.

**Figure 13.**
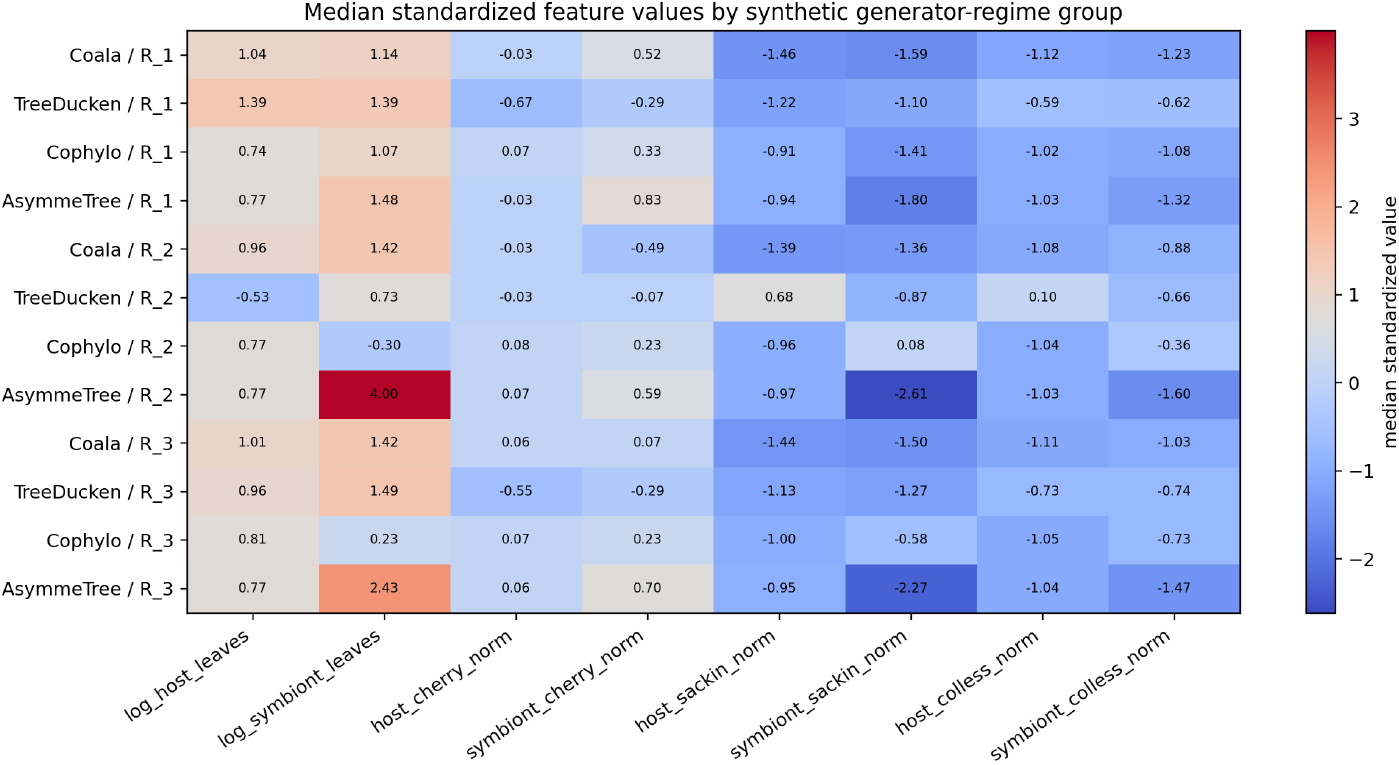
Median standardized feature values for each generator-regime combination. The features were standardized using the mean and standard deviation of the real datasets. A value of zero corresponds to the empirical mean. Positive values indicate values above the empirical mean, while negative values indicate values below it. Each value is expressed in empirical standard deviations.

The largest deviation from the empirical reference occurs for symbiont-tree size in `AsymmeTree` under *R*_2_, whose median standardized log-number of leaves is 4.00 standard deviations above the empirical mean. `AsymmeTree` also produces larger symbiont trees in *R*_1_ and *R*_3_. This large deviation likely contributes substantially to its low proximity and coverage in *R*_2_. The symbiont Sackin and Colless indices of `AsymmeTree` are also markedly below the empirical mean, particularly in *R*_2_ and *R*_3_.

For `cophylo`, symbiont-tree size varies more moderately across regimes. The median standardized log-number of symbiont leaves is above the empirical mean in *R*_1_ (1.07), slightly below it in *R*_2_ (−0.30), and close to the empirical mean in *R*_3_ (0.23). Its symbiont Sackin and Colless indices are generally below the empirical mean, with the Sackin index in *R*_2_ being close to zero.

`Coala` produces host and symbiont trees that are larger than the empirical mean, while its Sackin and Colless indices are consistently lower. This indicates lower global tree imbalance according to these two indices. A similar tendency appears in several `AsymmeTree` and `treeducken` combinations.

Overall, proximity and coverage highlight complementary properties of the generators. `Coala` produces datasets that are almost always close to at least one real dataset, but covers a more restricted part of the reference set. `treeducken` provides the broadest coverage and also shows high proximity in *R*_1_ and *R*_3_. `cophylo` combines high proximity with broad coverage across the three regimes, while `AsymmeTree` depends strongly on the regime and differs most markedly from the real reference set in *R*_2_.

These results concern tree size and tree shape. They measure agreement with the reference set in the eight-dimensional tree-feature space and should not be interpreted as a complete measure of biological realism.

## 4. Conclusion

This study shows that cophylogeny simulators are not interchangeable. Even when they are calibrated to represent the same high-level coevolutionary regime, the four generators considered here produce systematic differences in tree size and shape, association structure, and in their ability to reproduce the intended cospeciation and host-switch frequencies. These differences reflect the distinct modeling assumptions and simulation mechanisms implemented by each generator.

Importantly, these generator-specific patterns also propagate to downstream analyses. When the simulated datasets are analysed using the same parsimonious reconciliation procedure and the same event costs, the inferred event profiles and the number of optimal solutions can differ substantially across generators. This effect is particularly pronounced in the host-switch-dominated regime.

The comparison with real host-symbiont datasets further shows that proximity to real data and coverage of the empirical reference space capture complementary properties. `Coala` produces datasets that are very frequently close to at least one real dataset, but covers a more restricted part of the reference set. `treeducken` provides the broadest coverage, while `cophylo` combines high proximity with broad coverage across the three regimes. `AsymmeTree` is more strongly dependent on the simulation regime and differs most markedly from the real reference set in the host-switch-dominated regime.

These results should not be interpreted as identifying a “best” generator, nor as providing a complete measure of biological realism. Rather, they show that the choice of generator can introduce systematic structural biases into simulation-based studies. Synthetic benchmarks should therefore account explicitly for generator-specific effects, and, where possible, rely on multiple generators or on calibrations informed by empirical data. Future work could extend the present comparison to larger and more diverse collections of real host-symbiont systems and to additional biological and structural features, with the aim of designing more representative and comparable synthetic benchmark sets.

## Acknowledgements

Generative AI tools were used during manuscript preparation for language editing and to assist in refining code used to produce figures and L^A^T_E_X tables. They were not used to generate scientific results or conclusions. All AI-assisted content and code were reviewed and verified by the authors, who take full responsibility for the final manuscript.

## Data, script, code, and supplementary information availability

Scripts, code and the real datasets used in this study are available online (https://github.com/Gabbo240900/synthetic_cophylo).

## References

Alcala N, Jenkins T, Christe P, Vuilleumier S (2017). Host shift and cospeciation rate estimation from co-phylogenies. Ecology Letters 20, 1014–1024. 10.1111/ele.12799.

Balbuena JA, Míguez-Lozano R, Blasco-Costa I (2013). PACo: A Novel Procrustes Application to Cophylogenetic Analysis. PLoS ONE 8. Ed. by Corrie S. Moreau, e61048. 10.1371/journal.pone.0061048.

Bansal MS, Alm E, Kellis M (2012). Efficient algorithms for the reconciliation problem with gene duplication, horizontal transfer and loss. Bioinformatics 28, i283–i291. 10.1093/bioinformatics/bts225.

Baudet C, Donati B, Sinaimeri B, Crescenzi P, Gautier C, Matias C, Sagot MF (2014). Cophylogeny Reconstruction via an Approximate Bayesian Computation. Systematic Biology 64, 416–431. 10.1093/sysbio/syu129.

Charleston MA (2003). Recent results in cophylogeny mapping. Advances in Parasitology 54, 303–330. 10.1016/s0065-308x(03)54007-6.

Charleston M (1998). Jungles: a new solution to the host/parasite phylogeny reconciliation problem. Mathematical Biosciences 149, 191–223. 10.1016/S0025-5564(97)10012-8.

Charleston MA (2009). A new likelihood method for cophylogenetic analysis. URL: http://www.it.usyd.edu.au/research/tr/tr636.pdf.

Colless DH (1982). Review of: Phylogenetics: the theory and practice of phylogenetic systematics. Systematic Zoology 31, 100–104. 10.2307/2413420.

Conow C, Fielder D, Ovadia Y, Libeskind-Hadas R (2010). Jane: a new tool for the cophylogeny reconstruction problem. Algorithms for Molecular Biology 5. 10.1186/1748-7188-5-16.

Cruaud A, Rønsted N, Chantarasuwan B, Chou LS, Clement WL, Couloux A, Cousins B, Genson G, Harrison RD, Hanson PE, Hossaert-Mckey M, Jabbour-Zahab R, Jousselin E, Kerdelhué C, Kjellberg F, Lopez-Vaamonde C, Peebles J, Peng YQ, Pereira RAS, Schramm T, et al. (2012). An Extreme Case of Plant–Insect Codiversification: Figs and Fig-Pollinating Wasps. Systematic Biology 61, 1029–1047. 10.1093/sysbio/sys068.

Deng J, Yu F, Li HB, Gebiola M, Desdevises Y, Wu SA, Zhang YZ (2013). Cophylogenetic relationships between Anicetus parasitoids (Hymenoptera: Encyrtidae) and their scale insect hosts (Hemiptera: Coccidae). BMC Evolutionary Biology 13. 10.1186/1471-2148-13-275.

Di Palma G, Matias C, Sinaimeri B (2026). Similarities, Differences and Biases in Cophylogenetic Models for Host-Symbiont Coevolution. In: Comparative Genomics. Springer Nature Switzerland, pp. 213–233. 10.1007/978-3-032-26891-4_11.

Dismukes W, Heath TA (2021). treeducken: An R package for simulating cophylogenetic systems. Methods in Ecology and Evolution 12, 1358–1364. 10.1111/2041-210X.13625.

Donati B, Baudet C, Sinaimeri B, Crescenzi P, Sagot M (2015). Eucalypt: Efficient tree reconciliation enumerator. Algorithms for Molecular Biology 10, 3. 10.1186/s13015-014-0031-3.

Escudero M (2015). Phylogenetic congruence of parasitic smut fungi (Anthracoidea, Anthracoideaceae) and their host plants (Carex, Cyperaceae): Cospeciation or host-shift speciation? American Journal of Botany 102, 1108–1114. 10.3732/ajb.1500130.

Etherington GJ, Ring SM, Charleston MA, Dicks J, Rayward-Smith VJ, Roberts IN (2006). Tracing the origin and co-phylogeny of the caliciviruses. Journal of General Virology 87, 1229–1235. 10.1099/vir.0.81635-0.

Gómez-Acevedo S, Rico-Arce L, Delgado-Salinas A, Magallón S, Eguiarte LE (2010). Neotropical mutualism between Acacia and Pseudomyrmex: Phylogeny and divergence times. Molecular Phylogenetics and Evolution 56, 393–408. 10.1016/j.ympev.2010.03.018.

Hafner MS, Nadler SA (1988). Phylogenetic trees support the coevolution of parasites and their hosts. Nature 332, 258–259. 10.1038/332258a0.

Hendricks SA, Flannery ME, Spicer GS (2013). Cophylogeny of Quill Mites from the Genus Syringophilopsis (Acari: Syringophilidae) and their North American Passerine Hosts. Journal of Parasitology 99, 827–834. 10.1645/ge-2400.1.

Hughes J, Kennedy M, Johnson KP, Palma RL, Page RDM (2007). Multiple Cophylogenetic Analyses Reveal Frequent Cospeciation between Pelecaniform Birds and Pectinopygus Lice. Systematic Biology 56, 232–251. 10.1080/10635150701311370.

Hugot JP (1999). Primates and Their Pinworm Parasites: The Cameron Hypothesis Revisited. Systematic Biology 48, 523–546. 10.1080/106351599260120.

Hugot JP (2003). New Evidence for Hystricognath Rodent Monophyly from the Phylogeny of Their Pinworms. In: Tangled Trees: Phylogeny, Cospeciation, and Coevolution. Ed. by Roderic D. M. Page. University of Chicago Press, pp. 144–173.

Jackson AP, Charleston MA (2004). A Cophylogenetic Perspective of RNA–Virus Evolution. Molecular Biology and Evolution 21, 45–57. 10.1093/molbev/msg232.

Jansen G, Vepsäläinen K, Savolainen R (2011). A phylogenetic test of the parasite-host associations between Maculinea butterflies (Lepidoptera: Lycaenidae) and Myrmica ants (Hymenoptera: Formicidae). European Journal of Entomology 108, 53–62. 10.14411/eje.2011.007.

Kellner K, Fernández-Marín H, Ishak HD, Sen R, Linksvayer TA, Mueller UG (2013). Co-evolutionary patterns and diversification of ant–fungus associations in the asexual fungus-farming ant Mycocepurus smithii in Panama. Journal of Evolutionary Biology 26, 1353–1362. 10.1111/jeb.12140.

Kundu S, Bansal MS (2019). SaGePhy: an improved phylogenetic simulation framework for gene and subgene evolution. Bioinformatics 35, 3496–3498. 10.1093/bioinformatics/btz081.

Lei BR, Olival KJ (2014). Contrasting Patterns in Mammal–Bacteria Coevolution: Bartonella and Leptospira in Bats and Rodents. PLOS Neglected Tropical Diseases 8, 1–11. 10.1371/journal.pntd.0002738.

Martínez-Aquino A, Ceccarelli FS, Eguiarte LE, Vázquez-Domínguez E, de León GPP (2014). Do the Historical Biogeography and Evolutionary History of the Digenean Margotrema spp. across Central Mexico Mirror Those of Their Freshwater Fish Hosts (Goodeinae)? PLOS ONE 9, e101700. 10.1371/journal.pone.0101700.

McLeish MJ, van Noort S (2012). Codivergence and multiple host species use by fig wasp populations of the Ficus pollination mutualism. BMC Evolutionary Biology 12, 1. 10.1186/1471-2148-12-1.

Mendlová M, Desdevises Y, Civáňová K, Pariselle A, Šimková A (2012). Monogeneans of West African Cichlid Fish: Evolution and Cophylogenetic Interactions. PLoS ONE 7. Ed. by Dmitry A. Filatov, e37268. 10.1371/journal.pone.0037268.

Menet H, Daubin V, Tannier E (2022). Phylogenetic reconciliation. PLOS Computational Biology 18, e1010621. 10.1371/journal.pcbi.1010621.

Moreno MA, Holder MT, Sukumaran J (2024a). DendroPy 5. URL: https://jeetsukumaran.github.io/DendroPy/index.html.

Moreno MA, Holder MT, Sukumaran J (2024b). DendroPy 5: a mature Python library for phylogenetic computing. Journal of Open Source Software 9, 6943. 10.21105/joss.06943.

Murray EA, Carmichael AE, Heraty JM (2013). Ancient host shifts followed by host conservatism in a group of ant parasitoids. Proceedings of the Royal Society B: Biological Sciences 280, 20130495. 10.1098/rspb.2013.0495.

Newman MEJ (2003). Mixing patterns in networks. Physical Review E 67. 10.1103/physreve.67.026126.

Newman M (2018). Networks. Oxford University Press. 10.1093/oso/9780198805090.001.0001.

Page RDM (1994). Parallel phylogenies: reconstructing the history of host-parasite assemblages. Cladistics 10, 155–173. 10.1111/j.1096-0031.1994.tb00170.x.

Paterson A, Gray RD, Clayton DH, Moore J (1997). Host-parasite co-speciation, host switching, and missing the boat. In: Host-parasite evolution: General principles and avian models. Ed. by D. H. Clayton and J. Moore. Oxford: Oxford University Press, pp. 236–250.

Paterson AM, Palma RL, Gray RD (2003). Drowning on Arrival, Missing the Boat, and X-Events: How Likely Are Sorting Events? In: Tangled Trees: Phylogeny, Cospeciation, and Coevolution. Ed. by Roderic D. M. Page. University of Chicago Press, pp. 287–309.

Paterson AM, Poulin R (1999). Have chondracanthid copepods co-speciated with their teleost hosts? Systematic Parasitology 44, 79–85. 10.1023/a:1006255822947.

Pennington PM, Messenger LA, Reina J, Juárez JG, Lawrence GG, Dotson EM, Llewellyn MS, Cordón-Rosales C (2015). The Chagas disease domestic transmission cycle in Guatemala: Parasite-vector switches and lack of mitochondrial co-diversification between Triatoma dimidiata and Trypanosoma cruzi subpopulations suggest non-vectorial parasite dispersal across the Motagua valley. Acta Tropica 151, 80–87. 10.1016/j.actatropica.2015.07.014.

Poulin R (2010). Network analysis shining light on parasite ecology and diversity. Trends in Parasitology 26, 492–498. 10.1016/j.pt.2010.05.008.

Ramsden C, Holmes EC, Charleston MA (2009). Hantavirus Evolution in Relation to Its Rodent and Insectivore Hosts: No Evidence for Codivergence. Molecular Biology and Evolution 26, 143–153. 10.1093/molbev/msn234.

Refrégier G, Le Gac M, Jabbour F, Widmer A, Shykoff JA, Yockteng R, Hood ME, Giraud T (2008). Cophylogeny of the anther smut fungi and their caryophyllaceous hosts: Prevalence of host shifts and importance of delimiting parasite species for inferring cospeciation. BMC Evolutionary Biology 8. 10.1186/1471-2148-8-100.

Rosenblueth M, Sayavedra L, Sámano-Sánchez H, Roth A, Martínez-Romero E (2012). Evolutionary relationships of flavobacterial and enterobacterial endosymbionts with their scale insect hosts (Hemiptera: Coccoidea). Journal of Evolutionary Biology 25, 2357–2368. 10.1111/j.1420-9101.2012.02611.x.

Sackin MJ (1972). “Good” and “bad” phenograms. Systematic Biology 21, 225–226. 10.1093/sysbio/21.2.225.

Schaller D, Hellmuth M, Stadler PF (2022). AsymmeTree: A Flexible Python Package for the Simulation of Complex Gene Family Histories. Software 1, 276–298. 10.3390/software1030013.

Ŝimková A, Morand S, Jobet E, Gelnar M, Verneau O (2004). Molecular phylogeny of congeneric monogenean parasites (Dactylogyrus): A case of intrahost speciation. Evolution 58, 1001–1018. 10.1111/j.0014-3820.2004.tb00434.x.

Simões PM (2012). Diversity and dynamics of Wolbachia-host associations in arthropods from the Society archipelago, French Polynesia. PhD thesis. University of Lyon 1, France.

Simões PM, Mialdea G, Reiss D, Sagot MF, Charlat S (2011). Wolbachia detection: an assessment of standard PCR Protocols. Molecular Ecology Resources 11, 567–572. 10.1111/j.1755-0998.2010.02955.x.

Sinaimeri B, Urbini L, Sagot MF, Matias C (2023). Cophylogeny Reconstruction Allowing for Multiple Associations Through Approximate Bayesian Computation. Systematic Biology 72. Ed. by Adrian Paterson, 1370–1386. 10.1093/sysbio/syad058.

Stolzer ML, Lai H, Xu M, Sathaye D, Vernot B, Durand D (2012). Inferring duplications, losses, transfers and incomplete lineage sorting with nonbinary species trees. Bioinformatics 28, i409–i415. 10.1093/bioinformatics/bts386.

Tofigh A, Hallett M, Lagergren J (2011). Simultaneous Identification of Duplications and Lateral Gene Transfers. IEEE/ACM Trans. on Comput. Biol. Bioinf. 8, 517–535. 10.1109/TCBB.2010.14.

Vanhove MPM, Pariselle A, Van Steenberge M, Raeymaekers JAM, Hablützel PI, Gillardin C, Hellemans B, Breman FC, Koblmüller S, Sturmbauer C, Snoeks J, Volckaert FAM, Huyse T (2015). Hidden biodiversity in an ancient lake: phylogenetic congruence between Lake Tanganyika tropheine cichlids and their monogenean flatworm parasites. Scientific Reports 5. 10.1038/srep13669.

Viale E, Martinez-Sañudo I, Brown J, Simonato M, Girolami V, Squartini A, Bressan A, Faccoli M, Mazzon L (2015). Pattern of association between endemic Hawaiian fruit flies (Diptera, Tephritidae) and their symbiotic bacteria: Evidence of cospeciation events and proposal of “Candidatus Stammerula trupaneae”. Molecular Phylogenetics and Evolution 90, 67–79. 10.1016/j.ympev.2015.04.025.

Wang Y, Mary A, Sagot MF, Sinaimeri B (2020). Capybara: equivalence ClAss enumeration of coPhylogenY event-BAsed ReconciliAtions. Bioinformatics 36, 4197–4199. 10.1093/bioinformatics/btaa498.

Weckstein JD (2004). Biogeography Explains Cophylogenetic Patterns in Toucan Chewing Lice. Systematic Biology 53, 154–164. 10.1080/10635150490265085.

Zeng Y, Román-Palacios C (2025). Macroevolutionary Rates of Species Interactions: Approximate Bayesian Inference from Cophylogenies. 10.1101/2025.05.15.653894.

